# Association of cochlear outer hair cell - type II spiral ganglion afferents with protection from noise-induced hearing loss

**DOI:** 10.1101/2021.11.06.467554

**Authors:** Jennie M.E. Cederholm, Kristina E. Parley, Chamini J. Perera, Georg von Jonquieres, Jeremy L. Pinyon, Jean-Pierre Julien, David K. Ryugo, Allen F. Ryan, Gary D. Housley

**Affiliations:** Translational Neuroscience Facility and Department of Physiology, School of Medical Sciences, UNSW Sydney, Sydney, NSW 2052, Australia; Department of Psychiatry and Neuroscience, Laval University, CERVO Brain Research Centre, Quebec, Canada G1J2G3; Garvan Institute of Medical Research, Sydney, NSW 2010, Australia; School of Medical Sciences, UNSW Sydney, Sydney, NSW 2052, Australia; Department of Otolaryngology, Head, Neck & Skull Base Surgery, St Vincent’s Hospital, Sydney, NSW 2010, Australia; Departments of Surgery and Neurosciences, University of California San Diego, and Veterans Administration Medical Center, La Jolla, CA 92093, USA

**Author notes:** Corresponding author (GDH).

**Keywords:** peripherin type III intermediate filament, *Prph* knockout, contralateral suppression, medial olivocochlear efferents, cochlear amplifier, auditory neural circuit, distortion product otoacoustic emission, outer spiral bundle

## Abstract

The medial olivocochlear (MOC) efferent feedback circuit projecting to the cochlear outer hair cells (OHCs) confers protection from noise-induced hearing loss and is generally thought to be driven by inner hair cell (IHC) - type I spiral ganglion afferent (SGN) input. Knockout of the *Prph* gene (*Prph*KO) encoding the peripherin type III intermediate filament disrupted the OHC - type II SGN innervation and virtually eliminated MOC – mediated contralateral suppression from noise delivered to the opposite ear, measured as a reduction in cubic distortion product otoacoustic emissions. Electrical stimulation of the MOC pathway elicited contralateral suppression indistinguishable between wildtype (WT) and *Prph*KO mice, indicating that the loss of contralateral suppression was not due to disruption of the efferent arm of the circuit; IHC – type I SGN input was also normal, based on auditory brainstem responses. High-intensity, broadband noise (108 dB SPL, 1 hour) produced permanent hearing loss in *Prph*KO mice, but not in WT littermates. These findings associate OHC-type II input with MOC efferent - based otoprotection at loud sound levels.

## Introduction

Mammalian sound perception requires mechanoelectrical transduction by cochlear inner hair cells (IHCs), each discretely innervated by multiple type I spiral ganglion afferent neurons (type I SGN). The adjacent outer hair cells (OHCs) lend critical hearing sensitivity (∼40 dB, or 100-fold) (Ryan and Dallos, 1975), converting sound-evoked receptor potentials into electromechanical force, amplifying and shaping basilar membrane vibration as a ‘cochlear amplifier’ (Ashmore, 2019). OHC–specific electromotility can be directly measured as otoacoustic emissions using a sensitive microphone placed in the ear canal. The cochlear amplifier is subject to dynamic efferent neural feedback, which contributes broadly to hearing performance, including attention to sound, sound localization, hearing perception plasticity, improved hearing discrimination in noise, and protection of hair cells and their auditory synapses from acoustic overstimulation (reviewed by (Lopez-Poveda, 2018, Fuchs and Lauer, 2019)). The principal neural feedback circuit to the OHCs involves the contralateral medial olivocochlear (MOC) efferent neurons within the superior olivary complex - ventral nucleus of the trapezoid body in the brainstem. MOC axons form the crossed olivocochlear bundle (COCB), which passes across the floor of the fourth ventricle, projecting to the ipsilateral cochlea, to synapse with the OHCs (Brown, 2014, Weisz et al., 2012). Through this MOC projection, contralateral acoustic stimulation strongly inhibits the ipsilateral cochlear amplifier (‘contralateral suppression’) (Althen et al., 2012, Maison et al., 2012, Brown et al., 2013, Fuchs and Lauer, 2019). It is generally thought that the sensory input from the contralateral cochlea driving the MOC feedback circuit arises from the IHC - type I SGN (Brown et al., 2003, Maison et al., 2016). This position is primarily founded on studies in guinea pig and cat where single MOC efferent fibres show recruitment from low sound levels (20 dB SPL), with sharp tuning curves and characteristic frequencies similar to those of adjacent type I SGN afferents (Robertson and Gummer, 1985, Liberman and Brown, 1986). However, evidence is emerging that at high sound levels, where MOC efferent feedback confers protection from noise-induced hearing loss (Park et al., 2017), input from the OHC – type II SGN pathway may complement the IHC - type I SGN drive of the MOC efferent feedback circuit. Contralateral suppression was maintained following ouabain treatment of the cochlea that selectively ablated type I SGN, while leaving the type II SGN innervation of OHCs intact (Li et al., 2018). Further, type II SGN drive to the cochlear nucleus region known to connect with the MOC efferent arm of the circuit noise has recently been established for noise levels relevant to MOC efferent - mediated otoprotection (Weisz et al., 2021).

The type II SGN are a small, enigmatic subpopulation (∼ 5%) of unmyelinated afferents that exclusively innervate the OHCs (Brown et al., 1988a, Barclay et al., 2011). Their peripheral neurites project radially past the IHCs to form outer spiral bundles (OSB) that run basally, each branching to provide *en passant*, afferent synapses on multiple OHCs. This distributed receptive field overlaps with the frequency map of the cochlear amplifier, which spreads for ∼¼ octave towards the basal (high frequency) region, corresponding to the active region of the cochlear amplifier (Berglund and Ryugo, 1987, Jagger and Housley, 2003, Brown, 2014). This is also congruent to a basal shift in the MOC efferent activation, which overlaps with the cochlear amplifier region as well as the type II SGN afferent map as sound levels increase (Brown, 1994, Brown, 2014). The correspondence between the distribution of type II SGN afferent terminals and OHC-dependent active biomechanics prompted Kim (1986) to propose that type II SGN contribute to the afferent input for the MOC efferent innervation of the OHCs as a ‘closed-loop’ biomechanical gain control circuit. In support of this concept, in a prior study of mice null for the type III intermediate filament protein peripherin (*Prph*KO), we demonstrated disruption of OHC – type II SGN innervation and an associated loss of acoustically-evoked contralateral suppression of the quadratic distortion product otoacoustic emission (DPOAE) (Froud et al., 2015). A subsequent examination of the *Prph*KO mouse model challenged the morphological phenotype and concluded that the loss of contralateral suppression in these mice was probably due to failure of MOC efferent transmission to the OHCs (Maison et al., 2016). In the present study we provide a detailed characterization of the disruption of the type II SGN afferent innervation of the OHC in *Prph*KO mice and show that while the IHC - type I SGN afferent input and MOC efferent pathways remain normally active, there is an associated loss of otoprotection from acoustic overstimulation. These data support the concept that OHC – type II SGN input provides complementary drive of the cochlear amplifier feedback control circuit at high sound levels.

## Materials and Methods

### Animals

Male and female adult 129Sv/C57BL/6 wildtype (WT) and *Prph*-null (*Prph*KO) mice on the same background were used for this study. For acoustically – evoked contralateral suppression hearing studies, and noise-induced hearing loss studies, mice were anaesthetised using intraperitoneal injections (i.p.) of a ketamine (40 mg/kg) / xylazine (8 mg/kg) / acepromazine (0.5 mg/kg) (k/x/a) cocktail. Further k/x/a was administered (half the initial k/x/a dose mixed with the equivalent volume of 0.9% saline) as required to keep the mice anaesthetised throughout the experiment (typically every 30 min). For the electrically-evoked MOC stimulation experiments mice were anaesthetised with an initial k/x/a injection as above, then kept anaesthetised with 0.5-1% isoflurane supplemented with O_2_ throughout the experiment. The level of anaesthesia and the O_2_ saturation of the mice were monitored using a pulse oximetry system (MouseOx, STARR Life Sciences, SDR Scientific, Australia). Mice were kept on a heat pad for the duration of the experiment with their core body temperature clamped at 37° C by feedback control (Right Temp, Able Scientific, Perth, Australia). Ophthalmic ointment was applied to the eyes once the animal was anaesthetised to prevent corneal drying. At the completion of the studies, the animals were euthanised using pentobarbital (Virbac Australia, 100 mg/kg of body weight at 100 mg/ml, i.p.). Experiments were conducted according to UNSW Sydney Animal Care and Ethics Committee approved protocols, which conform to the Australian code for the care and use of animals for scientific purposes (NHMRC, 2013). The design and communication of the study align with the ARRIVE guidelines (Kilkenny et al., 2010).

### Genotyping

The *Prph*KO mouse model was established in 2001 using 129Sv strain embryonic stem cells bearing the peripherin knockout construct (exon 1 deletion) that were injected into C57BL/6 strain blastocysts (Lariviére et al. 2002). To produce heterozygous *Prph*KO mice, the generated chimeric mice were crossed with C57BL/6 mice. These heterozygous mice were used as breeders to provide littermates of identical genetic background. The *Prph*KO mouse line was established at UNSW in 2008. PCR-based genotyping utilised the following primer sets: Prph-5ꞌ-UTR-F: 5ꞌ GCT ATA AAG CCG CCC CGC ATC 3ꞌ; Prph-exon1-R: 5ꞌ AGGGCTGCGTTCTGCTGCTC 3ꞌ; LacZ-R: 5ꞌ GTC CTG GCC GTA ACC GAC CC 3ꞌ; Imaging of the PCR amplicons following agarose gel electrophoresis was used to distinguish WT (452 bp), KO (640 bp) and heterozygous (452 + 640 bp bands) mice. Homozygous knockout genotype validation was confirmed by the absence of peripherin immunolabelled cochlear type II SGN (after Froud et al. 2015), as below.

### Immunohistochemistry

Mouse cochleae were dissected and fixed by scali perfusion of 4 % paraformaldehyde, decalcified for 14 days in 8 % EDTA and then cryoprotected using 30% sucrose. The cochleae were mounted (Optimum Cutting Temperature (O.C.T) compound, Tissue-Tek, Sakura Finetek, Torrance, CA, USA) and cryosectioned at 50 µm. For wholemount preparations, cochleae were dissected into 2 - 4 pieces at the apical and basal turns. In both free-floating cryosections and the wholemounts, non-specific binding was blocked with 10 - 15 % normal goat or donkey serum, 1 % Triton X -100 in phosphate buffered saline (PBS) for 1 h at room temperature (RT). Sections were then immersed in primary antibodies: Neurofilament heavy polypeptide (NF200, Sigma, Cat# N4142, RRID: AB_477272; rabbit, 1:5000); C-terminal-binding protein 2 / RIBEYE ribbon synapse maker (Schmitz et al., 2000) (CtBP2, BD Bioscience, Cat# 612044, RRID: AB_399431; mouse, 1:500); Tubulin beta-3 chain (β-III tubulin) (TuJ1, Covance, Cat# MMS-435P, RRID: AB_2313773; mouse, 1:1000); Peripherin (Everest Biotech, Cat# EB12405, RRID: AB_2783842; goat, 1:1000; specificity validated in mouse by Froud et al. 2015), Parvalbumin alpha (Swant, Cat# PVG-213, RRID: AB_2650496; goat, 1:1000); Vesicular acetylcholine transporter (VAChT) (Phoenix Pharmaceuticals Inc., Cat# H-V007, RRID: AB_2315530; rabbit, 1:100) in 5 - 10 % normal goat or donkey serum, 0.1 % Triton X - 100 in PBS, overnight at RT. Sections were then washed in PBS and appropriate secondary antibody was applied overnight at RT (anti-rabbit IgG AlexaFluor 594, anti-mouse IgG AlexaFluor 488/594 (1:1000) or anti-goat AlexaFluor 488 (1:1000) (Molecular Probes)), 5 % normal goat or donkey serum in PBS. Following a further PBS wash, nuclear labelling was achieved by incubating the sections for 5 min in DAPI (4′,6-diamidino-2-phenylindole; 1:5000; Sigma). Two final washes in PBS were performed before the sections were mounted on glass slides using Vectashield (Vector Laboratories) and imaged on a Zeiss confocal microscope (Zeiss 710 NLO). Data were obtained from 1 – 3 organ of Corti z stacks (30 µm) analysed per animal, 3 animals per group. Data for the quantitative analysis of ribbon synapses was obtained using 30 µm stacks of confocal images; 63x oil immersion objective; mid-cochlear level, parsing the position of the CtBP2-labelled puncta within 2.5 µm bins relative to the equator line of the DAPI-labelled OHC nuclei. These data were blinded, randomized, and double scored before decoding.

### Hearing function tests

Hearing testing was carried out in a sound-attenuating chamber (Sonora Technology, Japan) using an auditory-evoked potential and DPOAE workstation (TDT system 3 with RX6 and RX6-2 signal processors, Tucker Davis Technologies, Ft Lauderdale, FL, USA) with BioSig32 software. Sound levels were calibrated using a one-quarter-inch Free Field Measure Calibration Microphone (model 7016; ACO Co Ltd., Japan).

### Acoustically-evoked contralateral suppression

The contralateral ear was exposed to broadband suppressor sound stimulation (96 dB SPL noise, 15 - 25 kHz, 15 s; n = 7 each for WT and *Prph*KO; 82 dB SPL noise, 10 - 17 kHz, 60 s; n = 8 each for WT and *Prph*KO) using a MF1 speaker (TDT), while cubic (2*f*_1_ -*f*_2_) DPOAEs were detected using a microphone coupled to the ipsilateral ear canal (ER-B10+, Etymotic Research, IL, USA), alongside two EC1 electrostatic speakers (TDT), controlled by the TDT system 3 workstation. DPOAEs were elicited using equal primary tones (*f*_1_ and *f*_2_; *f*_2_/*f*_1_ ratio: 1.25) around 20 kHz at 60 dB (for the 96 dB SPL noise study) or around 28 kHz at 65 dB (for the 82 dB SPL noise study). 10 - 50 measurements were averaged (6.7 / s) for each recording. DPOAE measurements were taken before (baseline), during and following (recovery) suppressor stimulus.

### Electrically-evoked contralateral suppression

To directly drive the contralateral MOC efferent innervation of the OHCs, these fibres were electrically stimulated at the point where they cross the midline on the floor of the fourth ventricle of the brainstem. Mice were initially anaesthetized with a k/x/a cocktail, maintained on a heat pad and eye ointment applied as described in the section *Animals*. This was followed by shaving the head to provide a clean surgical field for the dorsal-occipital approach to the fourth ventricle. A tracheostomy was performed, and the animals were ventilated throughout the experiment with a respiration rate of 100 breaths per minute and 10 cm peak inspiratory pressure (Kent Scientific TOPO™ Dual Mode Ventilator, Torrington, CT, USA). The animal was then positioned onto a teeth bar to stabilize the head position. The dorsal brainstem was accessed by removing muscles and connective tissue between the C1 (Atlas) vertebra and the occiput. This was followed by an incision in the dura covering the foramen, and the stimulating electrodes (two platinum iridium wires with 500 µm exposed tips, 400 µm separation) were positioned with one electrode on the midline of the fourth ventricle floor and the second electrode ipsilateral, using a micromanipulator. Twitching of the ears in response to a test electrical stimulation (monophasic 150 µs pulses, 200 Hz) ∼ 5s, indicated the correct position. α-D-tubocurarine (1.25 mg/kg, i.p.) was then administered to achieve muscle paralysis, which included paralysis of the stapedius reflex. The right (ipsilateral) external auditory meatus of the mouse was coupled to the DPOAE probe. A baseline measurement was obtained as the cubic DPOAE around 16 kHz at 50 or 55 dB SPL primaries (*f*_1_ = *f*_2_ amplitude, producing DPOAEs ∼ 10 dB above the noise floor); 20 averaged measurements (10 samples each) over 1 minute. The electrically-evoked contralateral suppression was then recorded during 25 seconds of stimulation (8 data point averages), followed by 2.5 min of recorded recovery. The DPOAE amplitudes were analysed relative to the noise floor. These experiments proved particularly challenging, with 3 / 16 WT mice and 3 / 7 KO mice providing useable data. However, electrically-evoked DPOAE suppression for each of the successful WT and *Prph*KO mice (n = 3 mice per genotype) were obtained from 3 – 5 repeats per mouse.

### Comparison of vulnerability to noise-induced hearing loss

To assess the otoprotection conferred by OHC - type II SGN – mediated MOC efferent cochlear amplifier suppression, noise-induced hearing loss was assessed in *Prph*KO versus WT mice using auditory brainstem responses (ABRs) and cubic DPOAE measurements with acute noise presentation. For ABR, following k/x/a anaesthesia induction, subdermal platinum electrodes were inserted subcutaneously at the vertex (+), over the mastoid process (-), and in the hind flank (ground) (after Cederholm et al., 2012). Click (100 µs alternating polarity) or pure tone pips (4, 8, 16, 24 and 32 kHz) stimuli (5 ms, 0.5 ms rise/fall time, 10/s) were delivered using an EC1 electrostatic speaker, with the generated ABR potentials amplified, filtered and averaged 512 times using the TDT System 3 workstation. The ABR threshold for each frequency was determined as the lowest intensity (5 dB steps from 70 dB SPL) at which the P2 ABR wave could still be visually observed above the noise floor (∼ 100 nV). Cochleae were then exposed to 108 dB SPL ‘open field’ broadband white noise (4 - 32 kHz; 2^nd^ order Bessel filter) for 1 hour. The white noise stimulus was generated using custom software with a National Instruments A/D driving an amplifier (BIEMA model Q250, Altronic, Northbridge, WA, Australia) and delivered via an MF1 speaker (TDT) positioned at midline, 15 cm in front of the mouse (n = 9 for each genotype). The noise was calibrated at ear level. ABR threshold shifts were determined immediately post-noise by remeasurement before the mice recovered from the anaesthesia. The mice were then rested for 14 days before being re-tested to determine irreversible hearing loss (permanent threshold shift (PTS)).

### Data analysis

Data are presented as the population mean ± S.E.M.. Statistical analysis was performed using a one-way Analysis of Variance (ANOVA), a two-way repeated measures (RM) ANOVA (Sigmaplot®, Systat Software Inc., San Jose, CA, USA), a two-way ANOVA or t-test, as indicated; significance at alpha ≤ 0.05. Data were tested for normal distribution and Holm-Sidak *post hoc* analysis was utilised for multiple pairwise comparisons within ANOVA. Grubbs’ test was used to assess data outliers (GraphPad software, San Diego, CA, USA).

## Results

### Disruption of the outer spiral bundle (type II SGN) fibre tracts in *Prph* knockout cochleae

The normal representation of type I and type II SGN neurite projections within the adult mouse cochlea was established in wildtype (WT) tissue using β-III tubulin and NF200 immunofluorescence (Figure 1; Video 1). The type II SGN sub-population was specifically identified using peripherin immunofluorescence in the WT cochleae (Figure 1A). The type II SGN somata were predominantly located in the lateral aspect of Rosenthal’s canal, juxtaposed to the intraganglionic spiral bundle of efferent fibres. The type II neurites track alongside the type I radial fibres within the osseous spiral lamina. Both types of afferent neurites pass through the habenula perforata and project towards the base of the inner hair cells (inner spiral plexus, ISP). However, the β-III tubulin and NF200 immunofluorescence labelled type II SGN neurites extended beyond the inner hair cells, with the fibres crossing the tunnel of Corti well beneath the extensive medial olivocochlear (efferent) bundle fibres (MOC) (Figures 1 – 4). The type II fibres then become basally projecting outer spiral fibres within OSB beneath each of the three rows of OHCs and their associated Deiters’ cells (DC). The outer spiral fibres periodically separate from the OSB, to ascend between the DC processes and make afferent synapses at the bases of OHCs (for example, Figures 1B,C, 2A).

**Figure 1:**
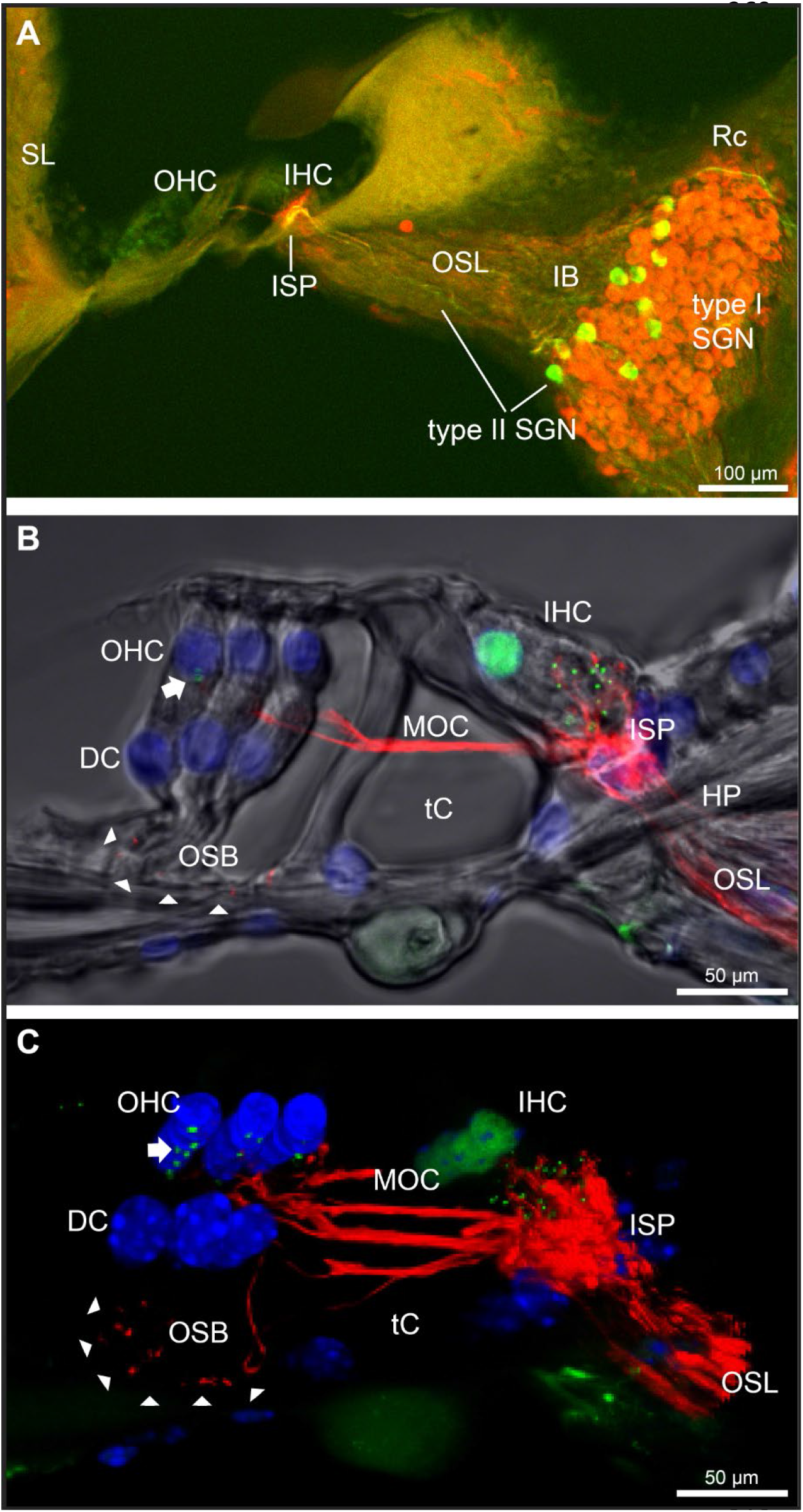
Immunofluorescence labelling of the afferent and efferent innervation of the organ of Corti in the wildtype mouse cochlea. **A**, Spiral ganglion neuron (SGN) somata and neurites (type I SGN (red) β-III tubulin immunofluorescence; type II SGN (green / yellow) peripherin immunofluorescence). The type II SGN sub-population is biased to the lateral aspect of Rosenthal’s canal (Rc), proximate to the intraganglionic spiral bundle (IB); the IB contains the medial olivocochlear (MOC) efferent axons from the superior olivary complex of the brainstem. All three nerve fibre types project to the organ of Corti via the osseous spiral lamina (OSL). The inner spiral plexus (ISP) is located at the basal pole region of the inner hair cells (IHC) and predominantly reflects type I SGN terminals. OHC, outer hair cells; SL, spiral ligament. **B,** Detail of the innervation and pre-synaptic ribbon complexes of the hair cells within the organ of Corti, delineated using NF200 (red) immunofluorescence for the nerve fibres and CtBP2/RIBEYE (green) immunofluorescence for the ribbons. Single confocal optical section overlaid with transmitted light image. NF200 immunolabelling delineates type II SGN neurites as discrete outer spiral bundle (OSB) fibre tracts (delineated by arrowheads) beneath the OHCs and their supporting Deiters’ cells (DC). Nuclei labelled with DAPI (blue). MOC efferent fibres cross the tunnel of the organ of Corti (tC) to innervate the OHCs. Note synaptic ribbons basal to the OHC nuclei (arrow). In comparison, the sub-nuclear domain of the IHCs contains many more CtBP2 immunopositive synaptic ribbons, each of which is aligned to a single type I SGN neurite terminal. Habenula perforata (HP). **C**, Confocal immunofluorescence reconstruction of a 50 µm cryosection delineates cell structure within the organ of Corti. The CtBP2 immunopositive synaptic ribbons are localized in a highly regular pattern beneath the mid-ventral aspect of the OHC nuclei (one or two per OHC; arrow), whereas each IHC contains > 10 CtBP2 puncta. The type II SGN outer spiral fibres within the OSB are clearly delineated (arrowheads) by the NF200 immunofluorescence ventral to the DCs. – see also Video 1 for 3D rendering.

**Video 1.**
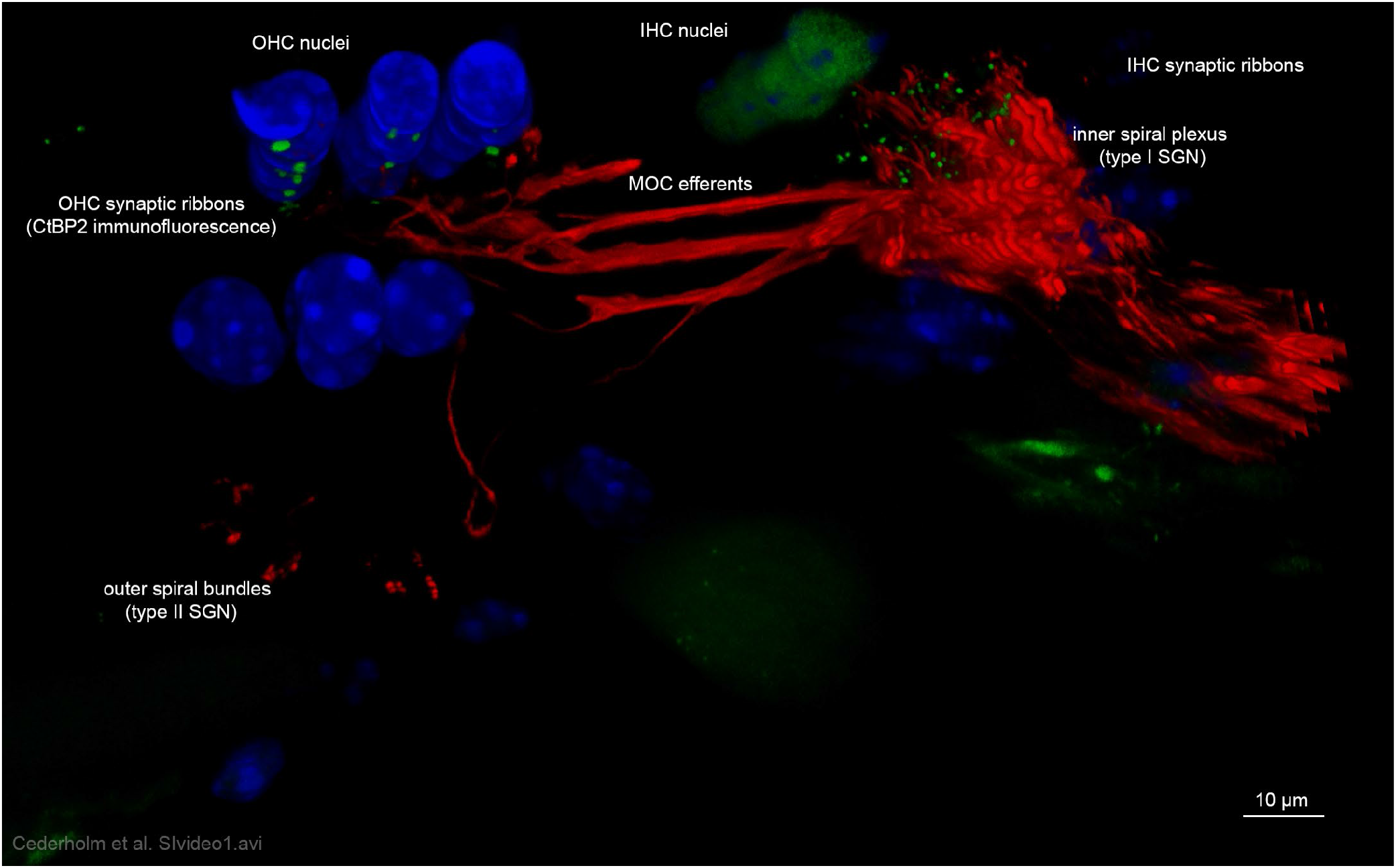
Projected confocal image stack of CtBP2 immunofluorescence (green) highlights the regular synaptic ribbon structure at the base of the three rows of outer hair cells, with one or two discrete puncta beneath the nucleus, compared with the dense synaptic ribbon complex associated with the basolateral membrane of each inner hair cell (WT cochlear organ of Corti). Type II spiral ganglion afferents (type II SGN) form the outer spiral bundle beneath the Deiters’ cells (NF 200 immunofluorescence – red) and project apically to the OHCs (upper left). This neural labelling includes the dense inner spiral plexus of the type I spiral ganglion neurite innervation (type I SGN) of each inner hair cell (IHC) – juxtaposed to the dense ribbon complex (upper right). The NF 200 immunofluorescence also extends to the dense matrix of medial olivocochlear (MOC) efferent fibres crossing the tunnel of Corti to innervate the outer hair cells. This projection extends the visualisation of Figure 1C. **Filename: Cederholm et al. SIvideo1.avi**

**Figure 2:**
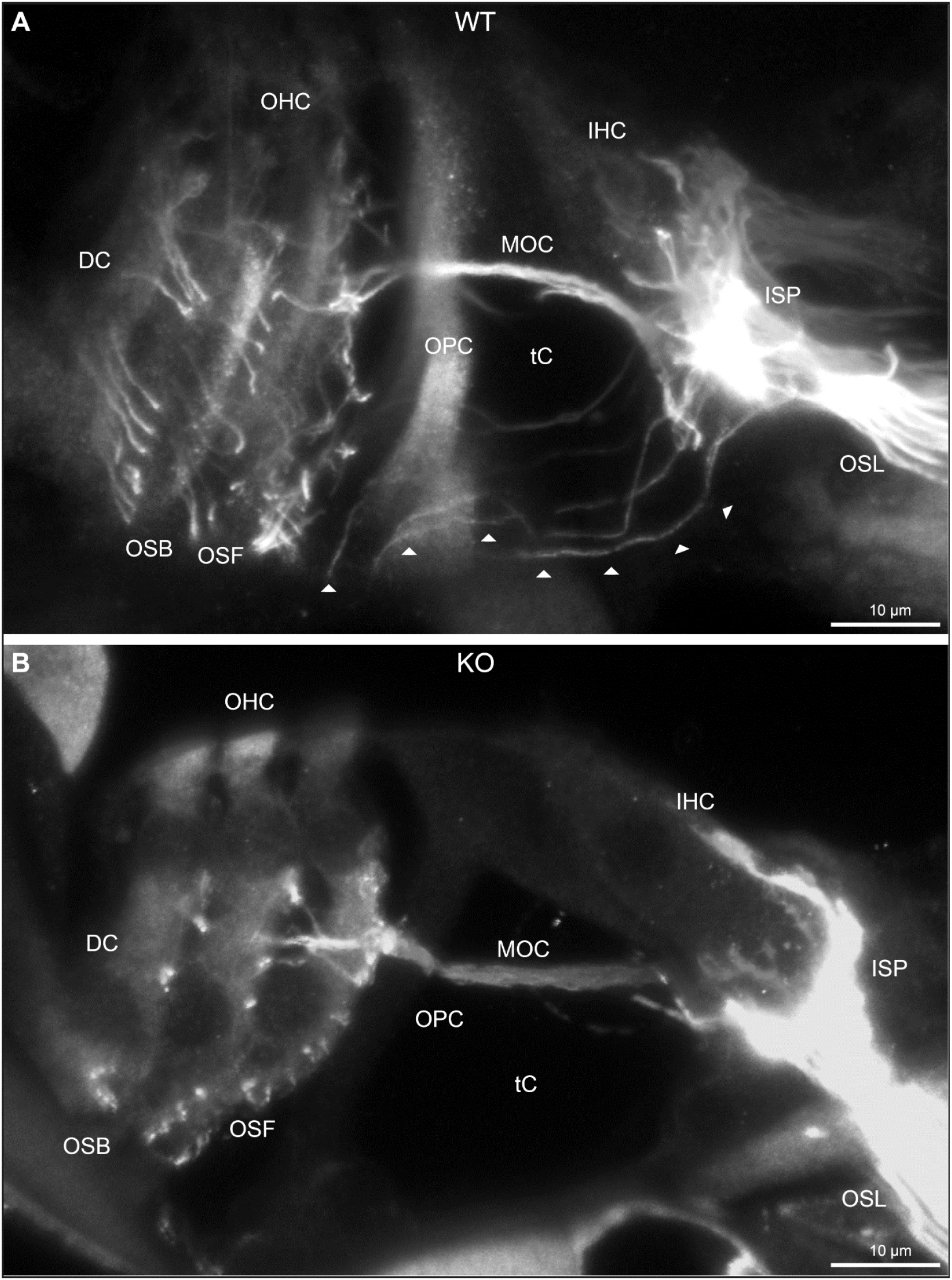
Disruption of type II spiral ganglion afferents, the outer spiral fibres (OSF), in the mid-basal region of the Prph knockout (KO) delineated using β-III tubulin immunofluorescence. **A**, In the WT cochlea, the type II spiral ganglion afferents exit from the basal aspect of the inner spiral plexus (ISP) beneath the inner hair cells (IHC), cross the floor of the tunnel of Corti (tC) (arrowheads), pass between the outer pillar cells (OPC) and turn basally to form parallel sheets of OSF along the sides of the Deiters’ cells (DC), as the outer spiral bundles (OSB). The OSF branch periodically and turn apically to terminate along the three rows of outer hair cells (OHC). The medial olivocochlear (MOC) efferent fibers cross the mid-region of the tC to innervate the OHC. **B**, In the PrphKO image, the OSF density is substantially reduced, while equivalent MOC efferent fiber projections extend to the OHC. OSL, osseous spiral lamina. Transparent-mode confocal reconstructed images of batch-processed WT and PrphKO cryosections; 21 – 25 µm depth z-stacks. See also Figure 2–figure supplement 1.

**Figure 3:**
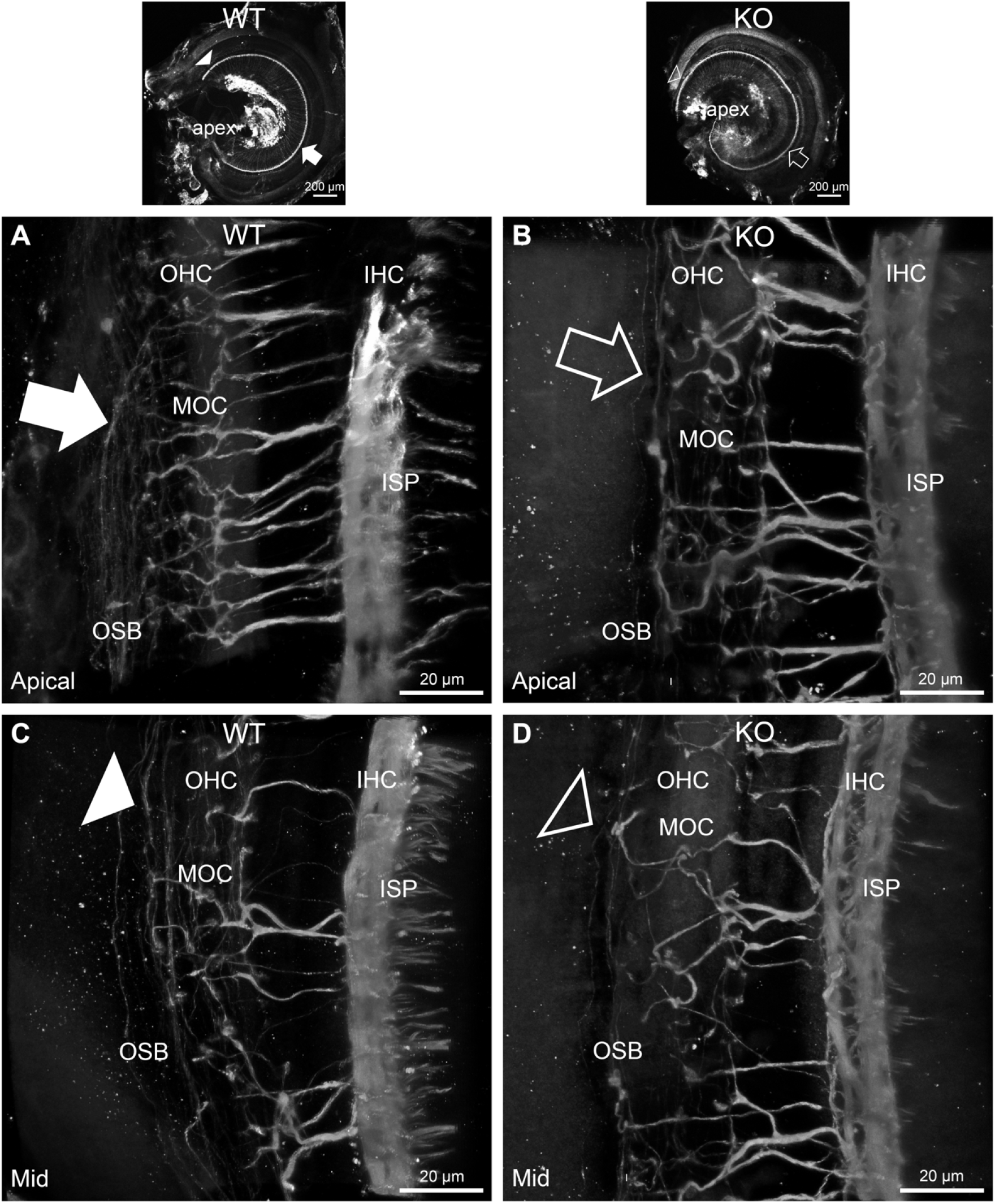
Disruption of the outer spiral bundle structure (OSB; type II spiral ganglion neurites form the outer spiral bundles) in Prph knockout (KO) vs. wildtype (WT) mouse cochleae from apex to mid region. Arrows in the low-power images indicate the corresponding regions of these whole-mount preparations which are shown at high resolution. The outer spiral fibres run parallel to the tunnel of Corti within the OSB (arrows), whereas the MOC fibres project across the tunnel from the inner spiral plexus (ISP) region to the outer hair cell (OHC) region, where they branch to innervate multiple OHCs. Note the comparatively higher number of outer spiral fibres in the apical region in the WT (filled arrow), (**A**), compared with the mid-cochlear region in the same tissue (filled arrowhead) (**C**). The corresponding regions in the wholemount KO cochlea (**B**, **D**; open arrow and open arrowhead) show minimal labelling in the OSB. Note the equivalency between KO and WT, of the medial olivocochlear (MOC) efferent axon projections to the OHCs. Maximum intensity projection β-III tubulin immunofluorescence confocal images of the organ of Corti (imaged from the basilar membrane surface to optimally resolve the OSB). IHC, inner hair cell region. – see also Videos 2 & 3 for 3D rendering.

**Video 2.**
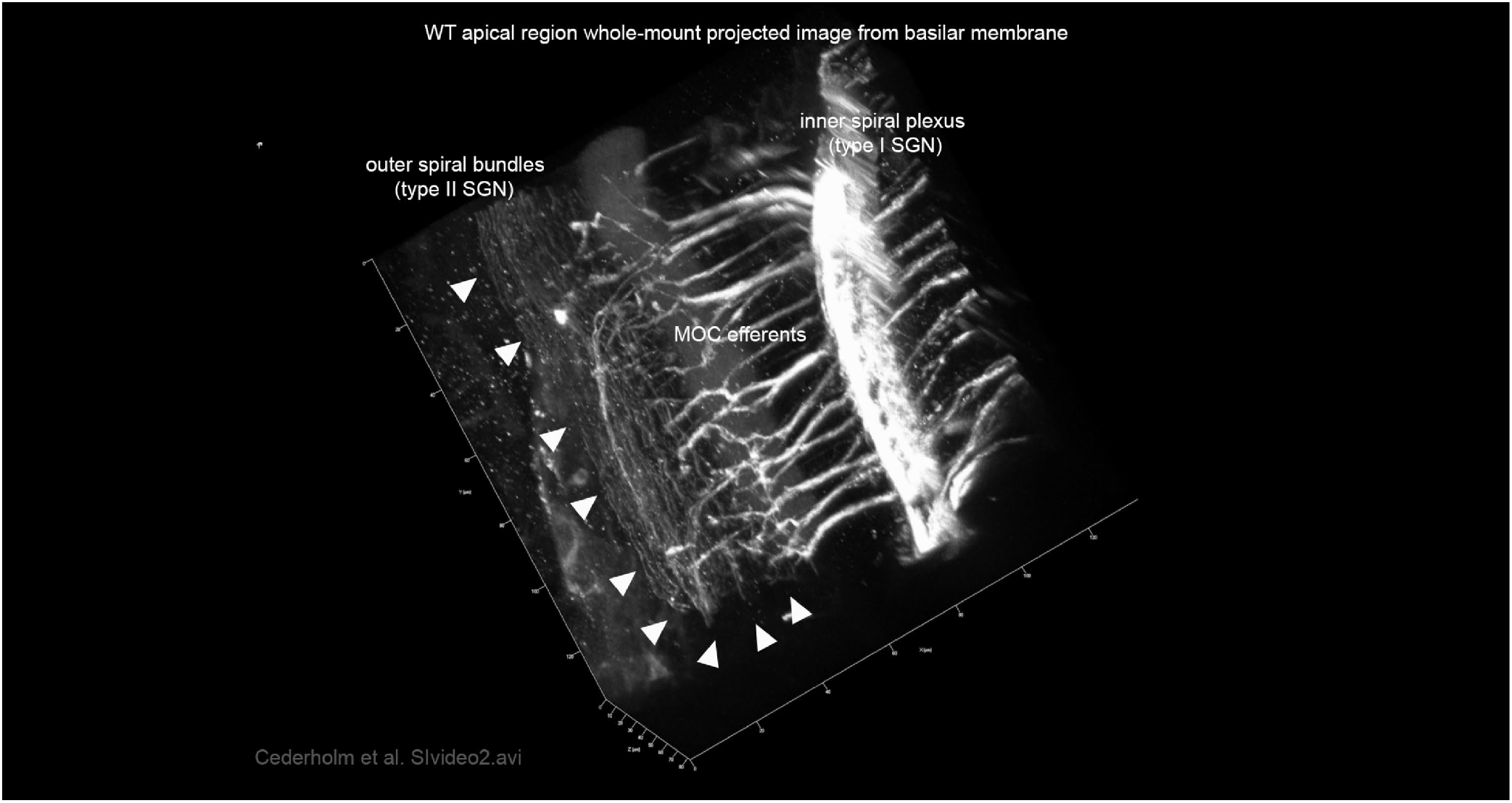
β-III tubulin immunofluorescence of innervation of the organ of Corti in the apical region of the wildtype mouse. Type II spiral ganglion afferents (type II SGN) form outer spiral bundles of small diameter unmyelinated neurites that run beneath the three rows of outer hair cells. Whole-mount maximum intensity projection of a confocal image stack (48 µm) imaged from the basilar membrane surface. The inner spiral plexus of type I spiral ganglion neurites (type I SGN) innervate the single row of inner hair cells. The medial olivocochlear (MOC) efferent fibres are shown crossing the tunnel of Corti to innervate the outer hair cells. This projection extends the visualisation of Figure 3A. **Filename: Cederholm et al. SIvideo2.avi**

**Video 3.**
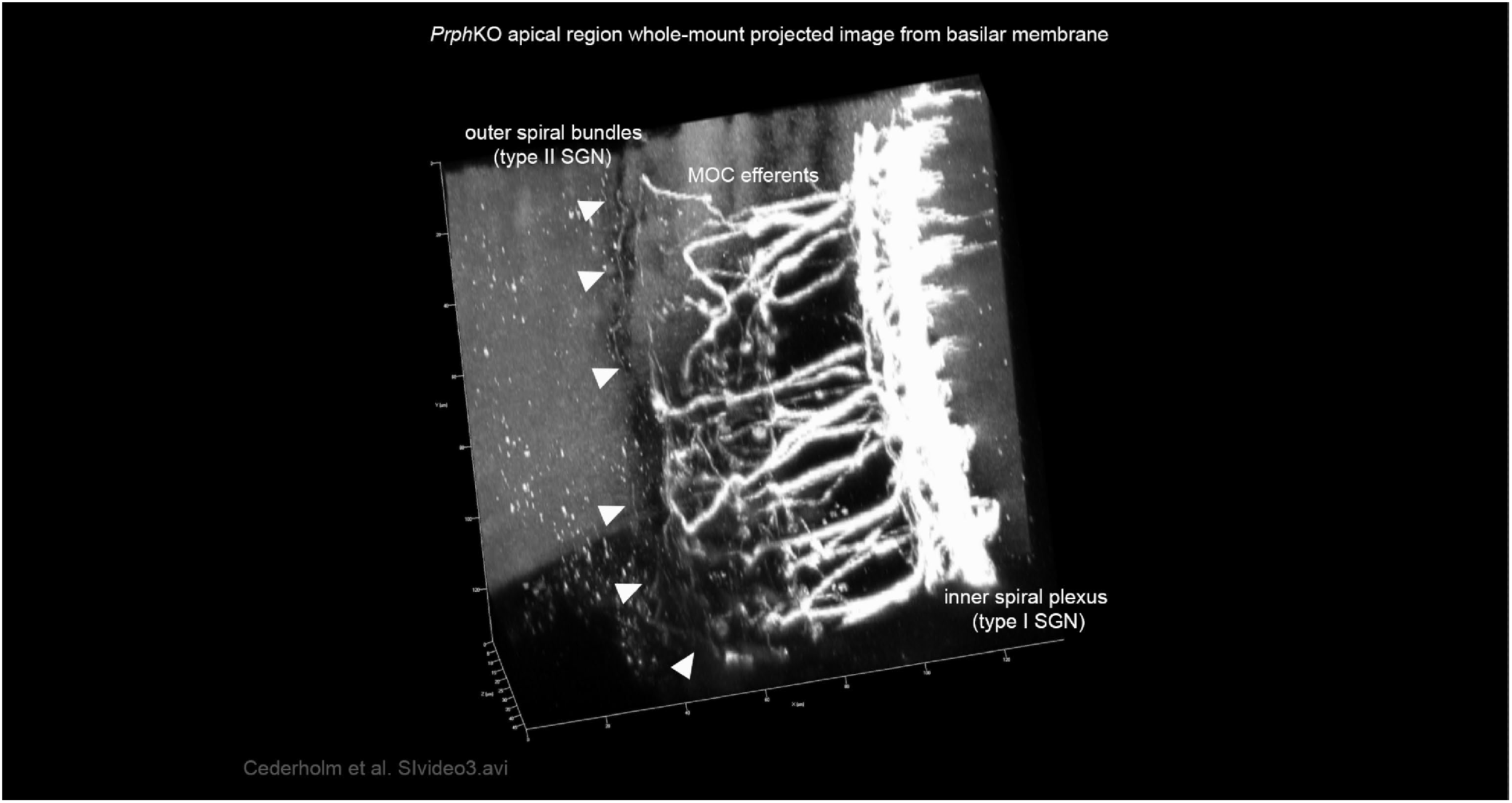
β-III tubulin immunofluorescence of innervation of the organ of Corti in the apical region of the PrphKO mouse. Whole-mount maximum intensity projection of a confocal image stack (48 µm) imaged from the basilar membrane surface from tissue batch-processed with the WT reference (SI Video 2). Note the reduced representation of Type II spiral ganglion afferents (type II SGN) in outer spiral bundles. Inner spiral plexus of type I spiral ganglion neurites (type I SGN) innervate the single row of inner hair cells. The medial olivocochlear (MOC) efferent fibres cross the tunnel of Corti to innervate the outer hair cells. This projection extends the visualisation of Figure 3B. **Filename: Cederholm et al. SIvideo3.avi**

**Figure 4:**
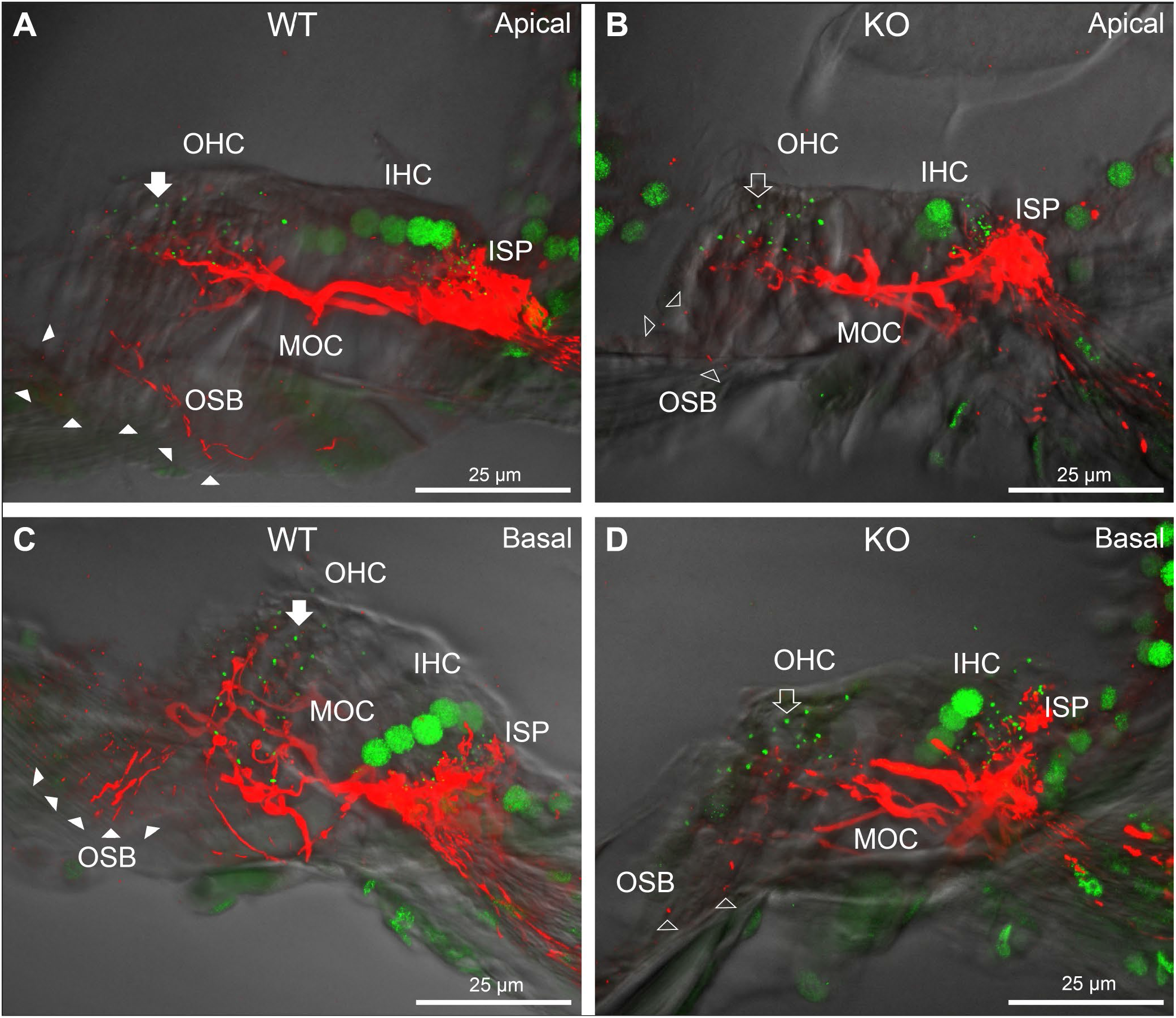
Comparison of NF200 (red) immunolabelling of outer spiral bundle (OSB) structure of the type II SGN neurites projecting to the outer hair cells (OHC) from wildtype (WT, filled arrow heads) and PrphKO (open arrow heads) cochleae. Note the limited OSB representation in the KO tissue from all regions. Images from single mid-modiolar cryosections for each of WT and KO illustrate the OSB fibre density within the organ of Corti at the apex and base. CtBP2/RIBEYE immunofluorescence (green puncta) delineates the pre-synaptic ribbons in both the OHCs and inner hair cells (IHC). The regular pattern of pre-synaptic ribbons at the mid-basal region of the three rows of OHCs evident in the WT (filled arrow), is disrupted in the KO (open arrow). The medial olivocochlear (MOC) efferent fibre innervation of the OHCs lies above the OSB. The dense type I SGN fibre synaptic complex at the base of the IHC (inner spiral plexus, ISP) is juxtaposed to the high density of synaptic ribbons in both WT and KO tissue. 50 µm cryosections; confocal projection images. See also Figure 4 –figure supplement 1

In *Prph*KO cochleae, the immunolabelling of the type II SGN afferent innervation was distinctly disrupted, whereas the type I SGN innervation and the MOC efferent fibre representation were normal in appearance. Comparisons between *PrphKO* tissue and corresponding WT tissue utilized batch immunoprocessing and imaging (Figures 2, – 4, Videos 2,3). In *Prph*KO, both β-III tubulin and NF200 labelling delineated a residual population of outer spiral fibres projecting from the ISP region, across the floor of the tunnel of Corti, and out to the OSBs in both whole-mount (n = 3 WT, n = 5 KO) and cryosectioned tissue (n = 6 WT, n = 4 KO). There was an evident gradient in this OSB pathology along the length of the organ of Corti, from an almost complete loss of fibres at the apex, to evident retention of some OSB structure in the basal region (Figure 4). It could not be established whether this reduction in outer spiral fibre number reflected atrophy of the peripheral neurite processes of the type II SGN, or arose from loss of type II SGN cell bodies, because the peripherin immunolabelling was intrinsically absent in the KO. However, the OSB remodelling clearly reflected disruption of the sensory neural drive from the OHCs.

Neither of the neuron-specific antibodies (β-III tubulin, NF200) delineated the type II SGN afferent terminal processes at the OHCs, likely reflecting limited trafficking of the β-III tubulin or the neurofilament 200 proteins to the synaptic region. Parvalbumin immunolabelling was therefore used to probe type II fibre terminations at the OHCs (Figure 5). Parvalbumin is a calcium binding protein that is expressed by type I and type II SGN, as well as the hair cells and supporting cells of the organ of Corti (Maison et al., 2016). Parvalbumin immunolabelling of the type II SGN fibres extended from the OSB underlying the Deiters’ cells, as outer spiral fibre projections to the synaptic pole of the OHCs. This further resolved the disruption of the OHC afferent innervation, where the density of these outer spiral fibres was substantially diminished in the KO tissue (n = 2), batch processed with WT tissue (n = 3).

**Figure 5:**
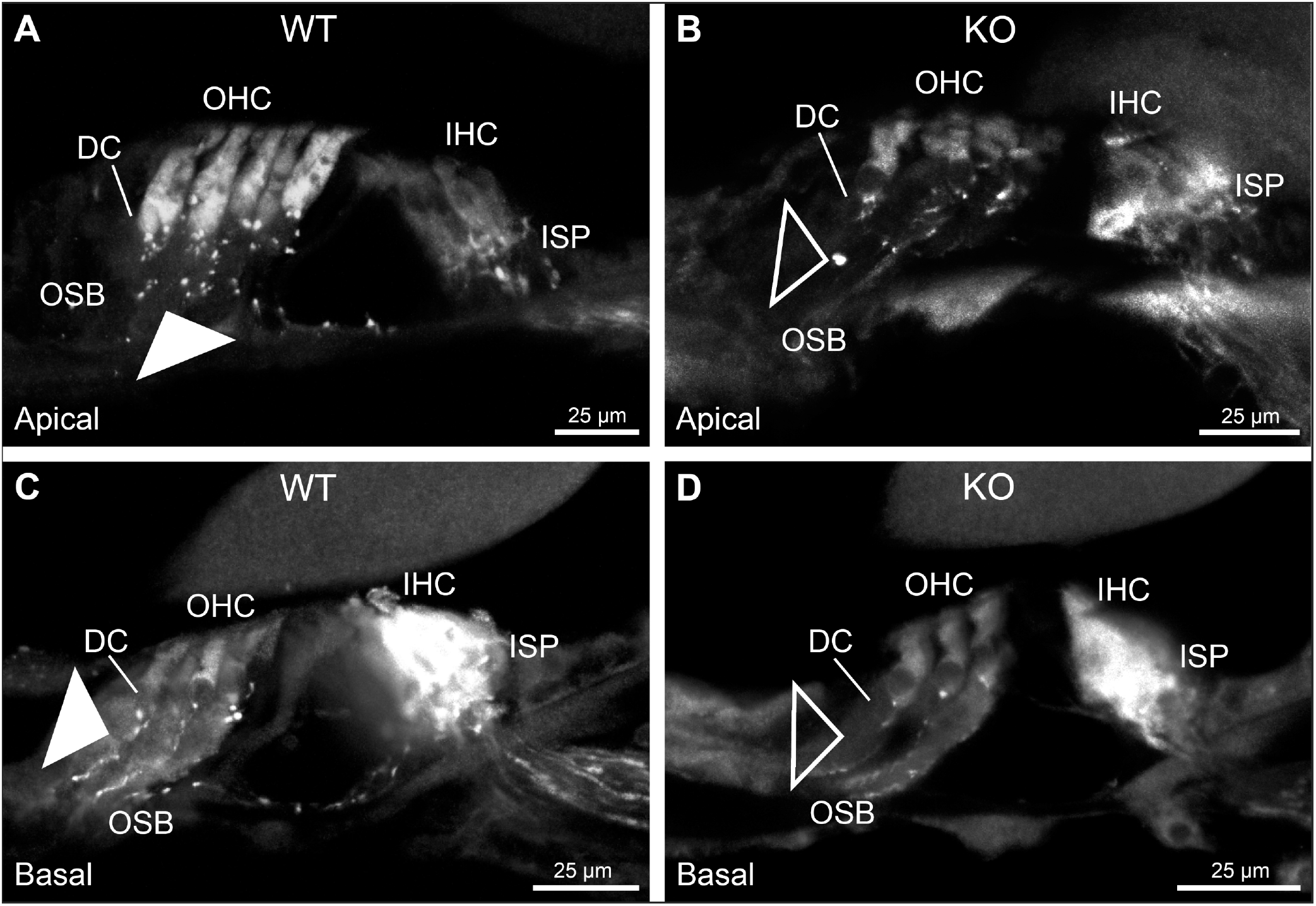
Parvalbumin immunofluorescence resolves the disruption of the type II SGN fibres (outer spiral fibres) within the outer spiral bundles (OSB) of the PrphKO mouse cochlea. Examples of batch-processed cryosections from WT (**A,C**) and KO (**B,D**) cochleae delineate the terminal regions of the outer spiral fibres, particularly with respect to their migration along the medial aspect of the Deiters’ cells (DC) up to the base of the outer hair cells (OHC). Parvalbumin was also strongly labelled in the inner hair cells (IHC) in the basal region of the cochlea (**C,D**), and in type I SGN afferent fibres within the inner spiral plexus (ISP). There was a much higher density of outer spiral fibre labelling at the level of the OSB in the WT tissue (filled arrowheads) compared with the KO tissue (open arrowheads), consistent with the β-III tubulin and NF200 immunolabelling experiments. The medial olivocochlear efferent projections to the OHCs are not delineated by this parvalbumin immunolabelling. Apical and basal regions from the same cochlea for each of WT and KO.

### Dysregulation of the outer hair cell pre-synaptic ribbon complexes in *Prph* knockout cochleae

CtBP2/RIBEYE immunolabelling of *Prph*KO cochleae showed disruption of the ordered structure of the synaptic ribbon complexes at the base of the OHCs, juxtaposed to the type II SGN afferent neurite terminals. As evident in Figure. 1C, Video 1, ∼ 2 CtBP2 positive puncta are normally present in the mid-basal region of each of the WT OHC, beneath the nucleus; in the KO tissue, this regular pattern was lost (Figure 6). This abnormality was elaborated by dual immunolabelling for CtBP2 and the vesicular acetylcholine transporter (VAChT), which marks the synaptic boutons of the MOC efferent innervation of the OHCs (Figure 6-figure supplement 1). In WT mice, multiple large VAChT positive efferent boutons are present at each OHC, on the medial and mid region of the basal pole of the cells. 1- 3 CtBP2-labelled pre-synaptic ribbons, corresponding to the type II afferent synapses, are interposed with the MOC synapses and solely occupy the lateral aspect of the synaptic pole of the OHCs. In the *Prph*KO cochleae, however, there was clear apical displacement of many of the synaptic ribbons relative to the base of the OHC, while the MOC efferent bouton distribution was unaffected, consistent with retention of the MOC innervation and disruption of the type II SGN innervation. This dislocation of the ribbon synapses was selective for the *Prph*KO OHCs, as CtBP2 and VAChT labelling of the IHC / ISP regions were comparable between WT and *Prph*KO cochleae (n = 5 WT, n = 3 KO). The displacement of the OHC pre-synaptic ribbon complexes in the *Prph*KO was quantified by batch-processing WT and *Prph*KO cryosections and undertaking a blinded measurement and analysis of the distance of the CtBP2 puncta relative to the equator line of OHC nuclei in z stacks of optical sections, with measurement of 13 – 57 OHC ribbons in cryosections from 5 WT mice (total 175 ribbons) and 6 – 60 ribbons from cryosections from 6 *Prph*KO mice (total 214 ribbons) (Figure 6, Figure 6-Source Data 1). Overall, the average CtBP2 puncta location in WT was 1.81 ± 0.24 µm below the OHC nucleus equator, compared with 0.53 ± 0.36 µm above the OHC nucleus equator for the *Prph*KO. The relative distribution of the ribbon complexes are compared in Figure 6C,D, resolving the significant apical migration of the ribbons in *Prph*KO OHCs. On average, across mice, the proportion of CtBP2 immunopositive synaptic ribbons located above the nucleus equator in WT OHC was 0.165 ± 0.054, whereas in the *Prph*KO mice, the apically located ribbons increased to an average proportion of 0.419 ± 0.036 (Figure 6D; *p* = 0.00313, t-test).

**Figure 6:**
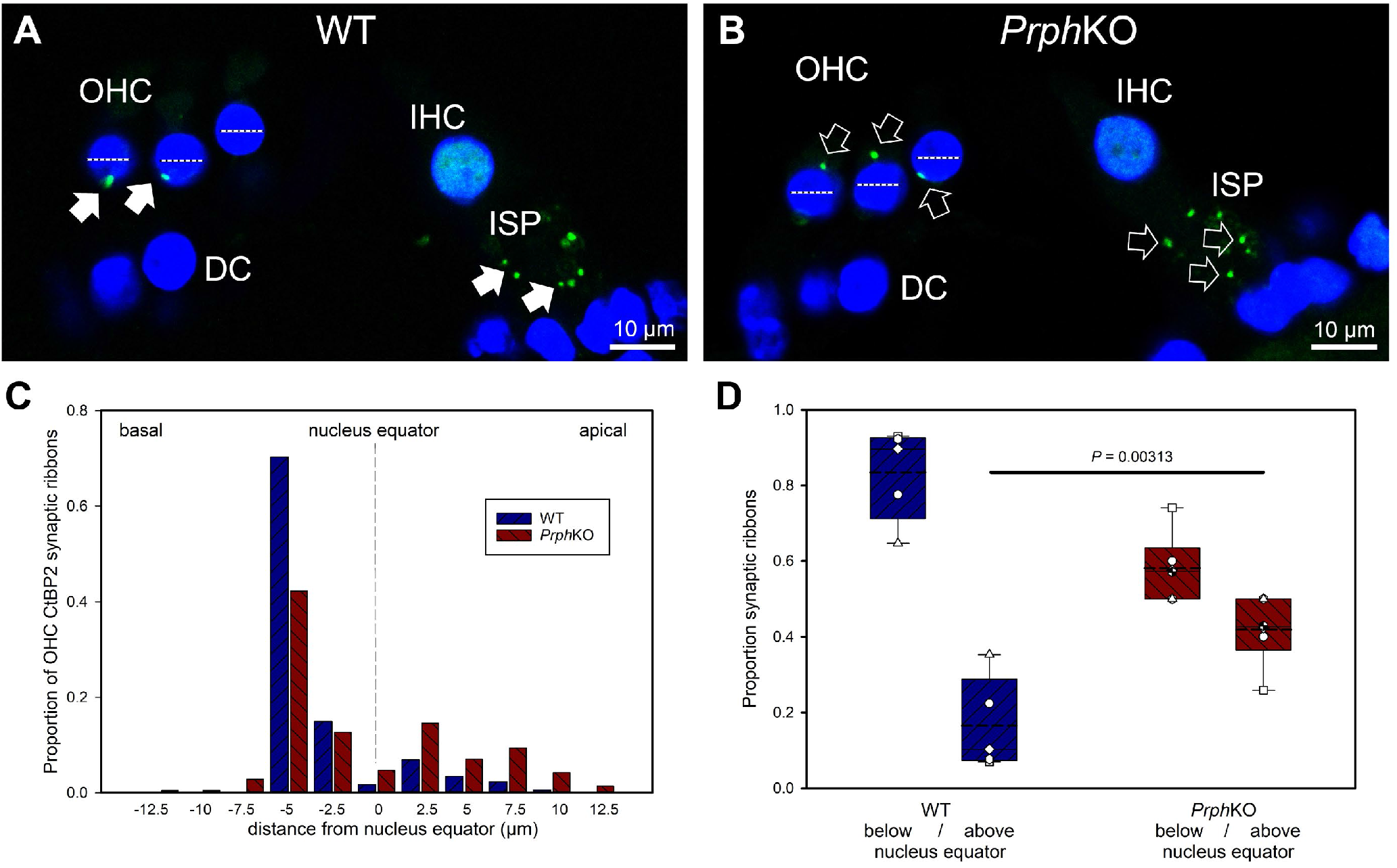
Quantitative analysis of the displacement of the outer hair cell (OHC) pre-synaptic ribbon complexes in PrphKO cochleae. **A**,**B** Examples of CtBP2/RIBEYE immunolabelling (green puncta, arrows) for WT and KO cochleae (50 µm cryosections; single confocal optical sections from Z stacks shown). Note the apical displacement of the OHC synaptic ribbons in the KO above the equator of the nucleus (dashed lines). DC, Deiters’ cells; ISP, inner spiral plexus; IHC, inner har cell; DAPI nuclear labelling (blue). **C**, Location of the synaptic ribbons relative to the OHC nuclei. Synaptic ribbon distribution from 175 WT OHC from 5 mice; 214 PrphKO OHC ribbons from 6 mice. The average number of ribbons per OHC was ∼ 2. **D**, Comparison of average distribution from the WT and KO mouse OHC ribbon complexes parsed as below or above the nucleus. The data indicate a significant migration of the synaptic ribbons away from the basal (synaptic) pole in the KO, consistent with loss of type II afferent fibre synapses (t-test). Box plots indicate 25% and 75% boundaries and 95% limits, with individual averages overlaid; dashed lines = means; solid lines = medians. See also Figure 6-figure supplement 1; Figure 6-Source Data 1.

### Acoustically-evoked contralateral suppression in the *Prph* knockout mouse

The phenotype of loss of contralateral suppression in the *Prph*KO mice reported by Froud et al. (Froud et al., 2015) was based on measurement of quadratic DPOAE (*f*_2_-*f*_1_) amplitude changes. This was validated in the current study using the cubic (2*f*_1_-*f*_2_) DPOAE (Fig. 10*A,B*), which is sensitive to changes in the gain of the cochlear amplifier (Mills and Rubel, 1996, Frank and Kossl, 1996), as mediated by MOC efferent drive. There was no difference in the baseline sensitivity of OHC electromechanical transduction in WT and *Prph*KO mice (reflected as equivalent cubic DPOAE amplitudes with 60 dB SPL *f*_1_ / *f*_2_ drivers). In WT mice, presentation of broad band white noise at 96 dB SPL (15 s) or 82 dB SPL (60 s) to the left ear produced robust reductions in the cubic DPOAE recorded in the right ear (contralateral suppression), which largely adapted within 15 s (Figure 7C,E). For 96 dB SPL contralateral noise, with 60 dB SPL pure tone drivers around 20 kHz in the ipsilateral ear, the peak noise-induced reduction in DPOAE in WT mice was -11.5 ± 3.0 dB (n = 7), measured 3 s after noise onset (Figure 7C). In contrast, only a residual contralateral suppression effect was detected in the KO mice at this level (peak = -1.6 ± 1.6 dB; *p* = 0.003, one sample t-test, n = 7). The difference in peak contralateral suppression between WT and KO mice with the 96 dB SPL noise presentation was therefore 9.9 dB (*p* = 0.013; unpaired t-test). The peak contralateral suppression in the WT mice with 82 dB SPL noise (65 dB SPL drivers around 28 kHz) was -3.5 ± 1.3 dB, measured at 6 s after noise onset (Figure 7D). There was no significant contralateral suppression in the KO mouse group at any sample period with 82 dB SPL noise presentation (e.g. -0.2 ± 0.1 dB at 6 s, *p* = 0.379). The 3.3 dB difference in peak DPOAE suppression at this noise level between KO and WT at 6 s was significant (*p* = 0.026, unpaired t-test; n = 8 per group).

**Figure 7:**
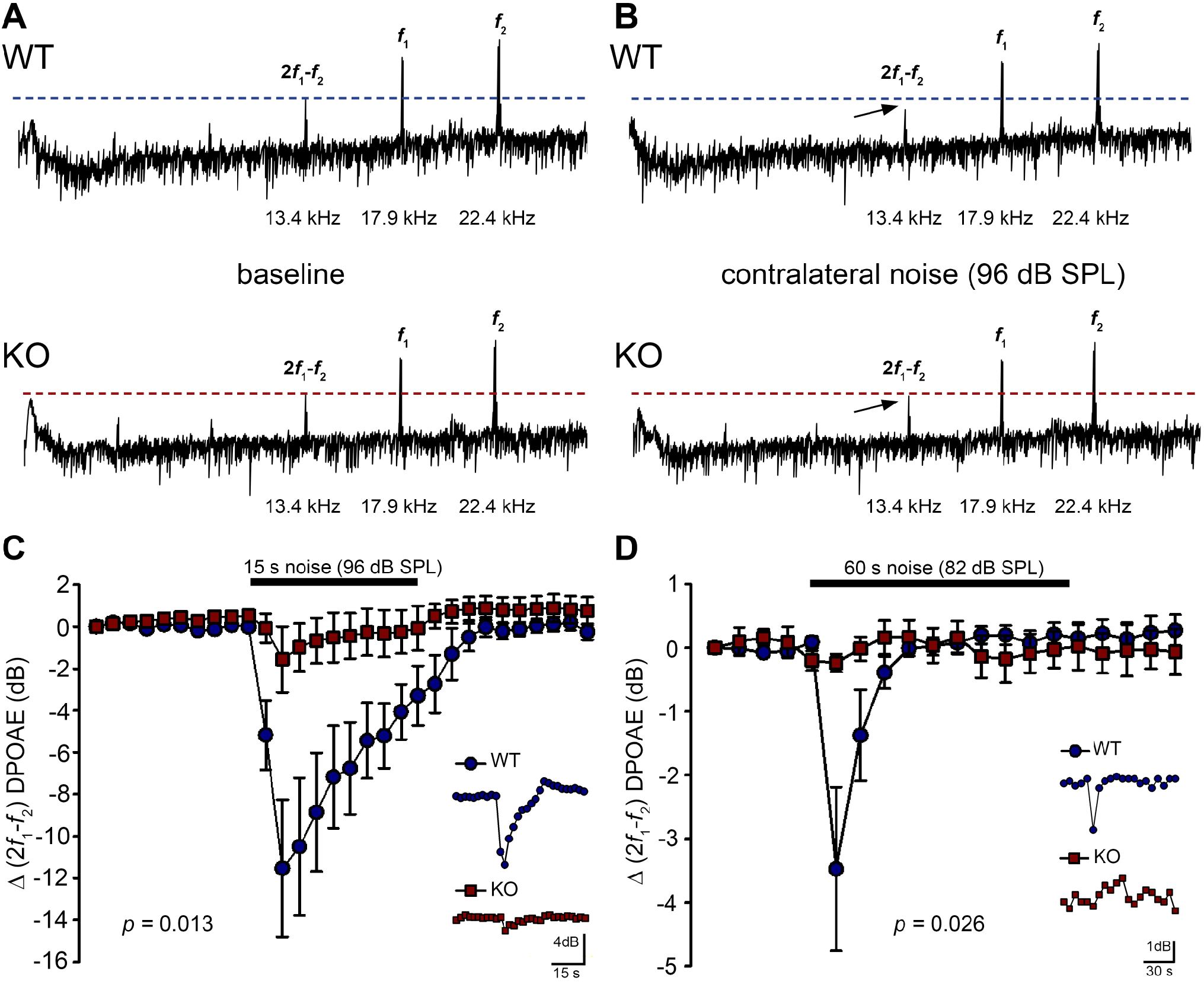
Contralateral suppression (CS) of the cubic (2f_1_-f_2_) DPOAE in wildtype (WT) and PrphKO mice under ketamine/xylazine/acepromazine anaesthesia. **A,B**, Examples of fast Fourier transforms of the DPOAE in the frequency domain with WT and KO at baseline (**A**) and (**B**) with the presentation of contralateral noise (96 dB SPL, 15 – 25 kHz, 15 s duration, with f_1_ and f_2_ about 20 kHz using 60 dB SPL drivers). The cubic DPOAE (arrows) is diminished in the WT mouse (contralateral suppression) but is not affected in the KO mouse. **C,** Data showing the difference in contralateral suppression with presentation of 96 dB SPL white noise in the WT mice and KO mice. Note the rapid onset to peak contralateral suppression at ∼ 3 s, followed by almost complete adaptation by 15 s. (unpaired t-test; n = 7 per group). **D,** Difference in contralateral suppression between WT and KO mice during 82 dB SPL noise (60 s, about 10 – 17 kHz, with f_1_ and f_2_ about 28 kHz using 65 dB SPL drivers). Note the complete adaptation by ∼ 30 s in the WT mice, whereas the DPOAE is unchanged in the KO mice (unpaired t-test; n = 8 per group).

### Validation of the viability of the medial olivocochlear efferent drive to the outer hair cells in the *Prph* knockout mouse by electrical stimulation

Direct electrical stimulation of the COCB efferent fibre track at the floor of the fourth ventricle was performed to eliminate the possibility that the loss of acoustically-evoked contralateral suppression in the *Prph*KO mice was due to disruption of the MOC efferent drive to the OHCs. Electrical stimulation of the MOC efferent fibres as they cross the floor of the fourth ventricle in the brainstem was used to directly compare the strength of the motor drive to the ipsilateral OHCs, assessed as cubic DPOAE suppression in WT versus *Prph*KO mice. This physiological preparation required stable presentation of DPOAE following a craniotomy to access the fourth ventricle, and precise micro-positioning of the bipolar stimulating electrodes, with one electrode at the mid-line, ∼ 0.2 mm rostral to the obex, and the second electrode positioned ∼ 0.4 mm lateral, towards the ipsilateral cochlea. Transient test stimuli were used with repositioning of the probe, until suppression of the cubic DPOAE was observed. The stimulus voltage was then lowered below threshold (∼ 3 V) and a series of recordings were obtained as the stimulus intensity was progressively stepped up to a maximum of 10 V, with four-minute recovery intervals between tests. The responses peaked within 12 s, with no substantial adaptation out to 25 s stimulation in both KO and WT mice. Figure 8A shows an example of electrically-evoked contralateral suppression obtained from a KO mouse. For statistical comparison, the amplitudes of electrically-evoked DPOAE suppression measurements at suprathreshold voltages were used (3 repeats for WT, 3 – 5 repeats for KO to determine the average response for each mouse) (Figure 8B). The average reduction in DPOAE amplitude during the 25 s stimulus from these trials was then used for statistical comparison between genotypes (Figure 8C). A repeated measures two-way ANOVA showed that the sensitivity of the electrically-evoked contralateral suppression was equivalent between KO and WT mice (WT = -2.7 ± 0.3 dB, n = 3; KO = -2.6 ± 0.2 dB, n = 3; *p* = 0.534). The need for precise positioning of the stimulus probe to achieve an electrically-evoked change in DPOAE is demonstrated in Figure 8A, where a repositioning of the probe ‘off-target’ eliminated the DPOAE suppression. There was no difference in the amplitudes of the baseline cubic DPOAE in the WT and *Prph*KO cochleae, WT average = 17.1 ± 0.3 dB above noise-floor; KO = 16.1 ± 0.2 dB; *p* = 0.078; unpaired t-test).

**Figure 8.**
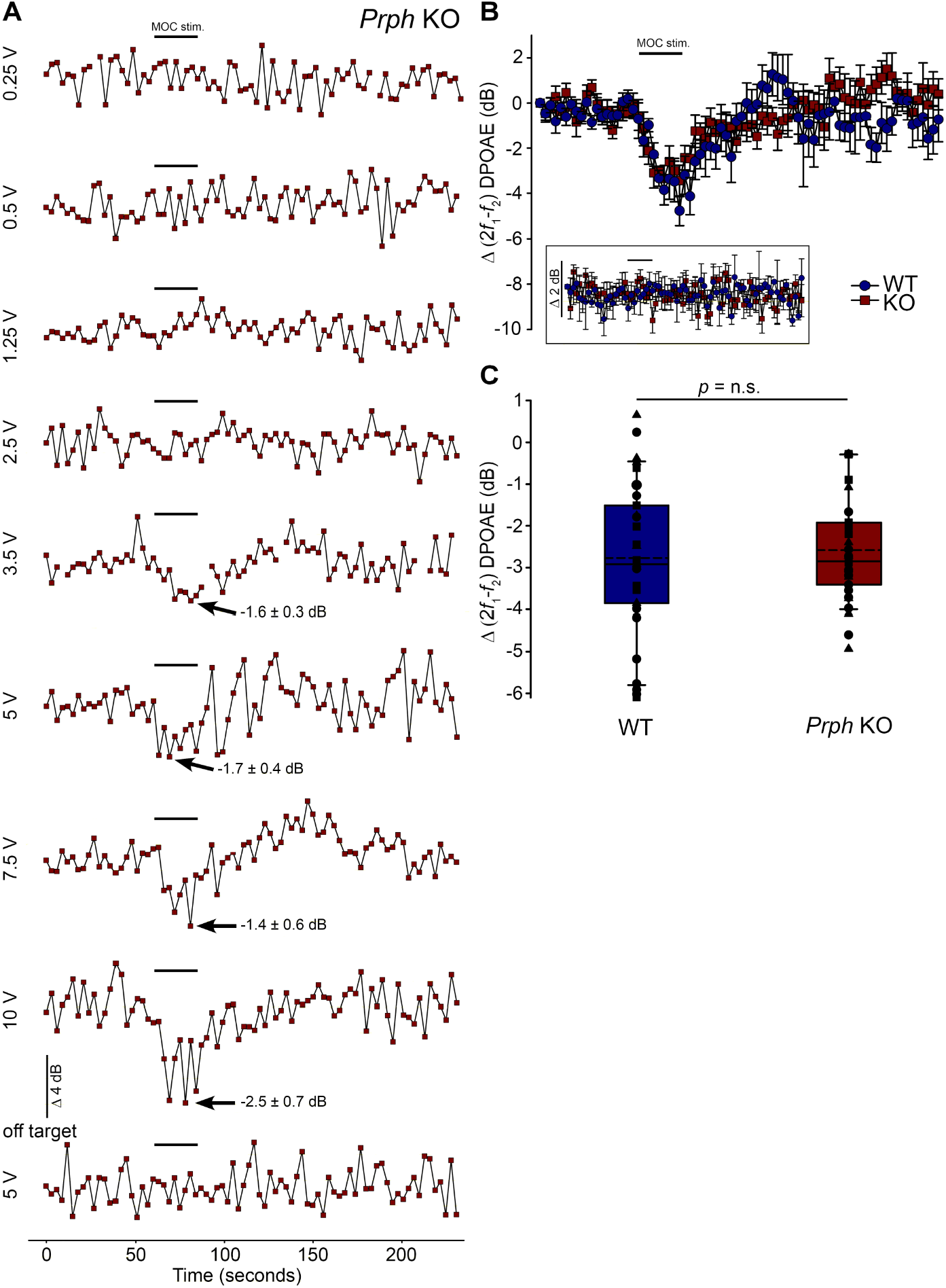
Equivalency of electrically-evoked DPOAE suppression in PrphKO and wildtype (WT) mice under isoflurane anaesthesia. **A**, Baseline cubic DPOAE (2f_1_-f_2_) with f_1_ and f_2_ primaries around 16 kHz at 50 - 55 dB SPL was established using 3 s repeated sampling for 60 s, followed by 25 s of direct electrical stimulation of the crossed olivocochlear bundle fibres on the floor of the brainstem fourth ventricle (medial olivocochlear efferent stimulation (MOC stim.), black bar; monophasic pulses, 150 µs duration, 200 Hz), and then 150 s of recovery (PrphKO mouse). The bottom trace (‘off-target’) illustrates a ‘false-negative’ outcome (a 2 mm rostral repositioning of the probe eliminated the suppression of DPOAE elicited with an equivalent 5 V stimulus level). Arrows indicate average contralateral suppression over the stimulus period relative to the preceding baseline average. **B**, Overlay of the averaged electrically-evoked DPOAE suppression responses for WT and KO mice. The data show the mean and S.E.M. based on 3 – 5 repeated MOC stimuli for each mouse, with 25 s electrical stimulation at suprathreshold stimulus voltages (WT 5V, n = 3; KO 2 – 8 V, n = 3). The inset shows stability of the recordings with subthreshold stimulation (0.25 – 1V). **C**, Boxplots show the equivalency of the average electrically-evoked DPOAE suppressions, with 25% and 75% boundaries and bars to 95% confidence limits. Individual data overlaid (symbols show repeats for individual mice); dashed lines show the mean; solid lines show the median. p = n.s. indicates p > 0.05, by repeated measure two-way ANOVA.

### Loss of otoprotection from noise-induced hearing loss in *Prph* knockout mice

Reduction of OHC electro-mechanical transduction by the MOC efferent pathway is otoprotective (Park et al., 2017, Housley et al., 2020). For example, surgical ablation of the COCB at the floor of the fourth ventricle in cats increased noise-induced threshold shifts (Rajan, 2000) and over-expression of the OHC α9 nicotinic acetylcholine receptor subunit in mice confers protection to NIHL (Maison et al., 2002). Here we investigated the effect of disruption of the OHC – type II SGN sensory input arising from knockout of *Prph* in this key aspect of hearing homeostasis. The vulnerability to 1 hour of acute white noise (4 – 32 kHz) at 108 dB SPL was assessed in WT and *Prph*KO mice by ABR (Figure 9). WT and KO mice had equivalent pre-noise ABR thresholds for click and tone-pip stimuli (4 – 32 kHz) – (Figure 9A), which indicates that disruption of the type II SGN sensory drive does not affect hearing sensitivity in a nominally quiet environment. ABR threshold shifts immediately after noise exposure were comparable between the WT and *Prph*KO mice, with the greatest changes (∼ 60 - 70 dB) observed from 16 – 32 kHz (Figure 9B). Permanent hearing loss (permanent threshold shift – PTS), was determined by re-measurement of ABR thresholds two weeks after noise exposure. While the WT mice did not exhibit any significant PTS, with thresholds returning to pre-noise baseline, the *Prph*KO mice exhibited high frequency hearing loss, with PTS of 10.9 ± 2.0 dB at 24 kHz and 19.4 ± 5.1 dB at 32 kHz (*p* < 0.016; Two-way RM ANOVA; n = 9 per genotype (Figure 9C). Data distribution from individual animals at 24 kHz and 32 kHz shown in boxplots (Figure 9C-figure supplement 1). The Power of the two-way ANOVA analysis of genotype effect on noise-induced hearing loss was 0.655 with alpha = 0.05.

**Figure 9.**
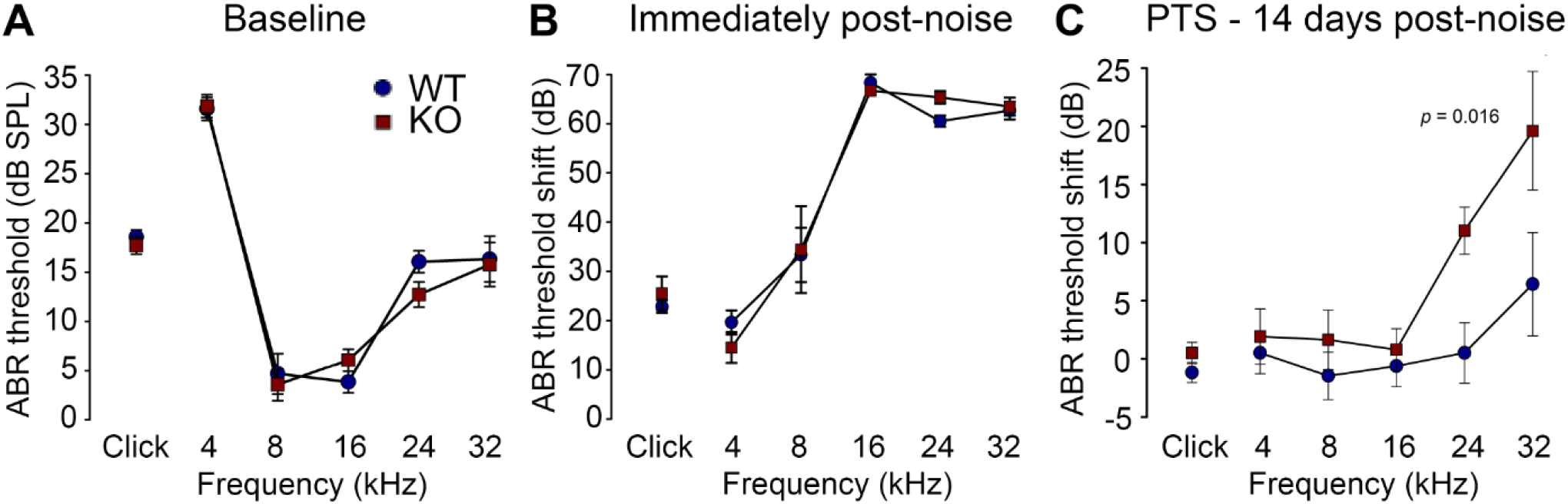
PrphKO mice are vulnerable to noise-induced high frequency hearing loss. **A**, Hearing sensitivity was equivalent in KO and wildtype (WT) mice based on auditory brianstem response (ABR) thresholds; SPL sound pressure level. (n = 9 per group) **B,** There was no significant difference in ABR threshold shifts immediately following exposure to 1 hour of 108 dB SPL white noise (4 – 32 kHz; open-field) between WT and KO mice. **C,** Permanent threshold shifts were determined two weeks post-noise. While WT mouse thresholds returned to pre-noise levels, KO littermates had significant permanent threshold shifts at frequencies above 16 kHz (ANOVA). Grubbs’ test (GraphPad) excluded one data point as an outlier for PrphKO 24 kHz. Ketamine/xylazine/acepromazine anaesthesia. (see also Figure 9-figure supplement 1; Figure 9-Source Data 1; Figure 9-figure supplement 2).

The ABR input / output functions for neural recruitment of the WT mice, and *Prph*KO mice 2 weeks post-noise, demonstrated equivalency at frequencies where the *Prph*KO mice exhibited significant threshold shifts (Figure 9-figure supplement 2). The average slopes for 24 kHz were: WT = 0.032 ± 0.002 µV/ dB SPL; *Prph*KO = 0.038 ± 0.005 µV/ dB SPL. 32 kHz slopes were: WT = 0.032 ± 0.003 µV/ dB; *Prph*KO = 0.038 ± 0.005 µV/ dB SPL (two-way RM ANOVA for genotype comparison, *p* > 0.05). Supporting comparable ABR input / output functions, cubic DPOAE (8 kHz, 12 kHz, 16 kHz, 24 kHz, 32 kHz) at two weeks post-noise showed no significant differences in thresholds (respective difference of means: 2.22 dB, 1.11 dB, 4.44 dB, 5.00 dB, 8.33 dB; *p* > 0.05 two-way RM ANOVA; input / output functions (*p* > 0.05 two-way RM ANOVA); consistent with synaptopathy underlying the ABR PTS.

## Discussion

These studies provide strong evidence that at loud sound levels, the OHC – type II SGN afferent input to the cochlear nucleus contributes to the activation of the MOC efferent feedback control of the cochlear amplifier to confer protection from noise-induced hearing loss. Hearing sensitivity (threshold ABR and DPOAE amplitude) is equivalent for WT and *Prph* deficient mice. However, in the *Prph*KO, acoustically-evoked contralateral suppression was absent at 82 dB SPL noise (just above the threshold for detectable WT contralateral suppression), and was ∼ 10 dB less than WT using high level (96 dB SPL) noise (where 3 dB represents a halving of intensity). These measurements were undertaken at 20 kHz – 28 kHz, probing the middle region of the mouse cochlea (Muller et al., 2005). The region exhibits profound SGN dendrite dysmorphology in the *Prph*KO cochlea (Fig. 3B), with only residual outer spiral fibres in evidence, and OHC synaptic ribbons displaced.

Disruption of type II SGN afferent structures in *Prph*KOs is consistent with evidence that peripherin supports axon growth, particularly in sensory fibers of the peripheral nervous system; e.g. in the dorsal root ganglia (DRG) (Helfand et al., 2003, Lariviere and Julien, 2004). Peripherin gene deletion similarly causes selective loss of small unmyelinated DRG fibres (Lariviere et al., 2002). In the mouse cochlea, peripherin is strongly biased to the unmyelinated type II SGN, over type I SGN from birth onward, and most strongly during early post-natal neurite extension and synaptic consolidation (Huang et al., 2012, Huang et al., 2007, Elliott et al., 2021). Spiral ganglion explants from the neonatal *Prph*KO mouse exhibit enhanced neurite outgrowth, suggesting altered response to guidance and/or stop cues (Barclay et al., 2010).

Our initial NF200 immunolabelling in adult *Prph*KO cochlea revealed significant reduction in the number of outer spiral fibres (Froud et al., 2015). In that study serial blockface scanning electron microscopy in WT and *Prph*KO mouse cochleae (Froud et al. 2015, Suppl. Fig. 2 and Suppl. Table 1) showed no type II SGN afferent boutons on OHCs in *Prph*KO organ of Corti, compared to ∼ 1 bouton per WT OHC, consistent with other mouse data (Berglund and Ryugo, 1987, Wood et al., 2021). These findings correlated with a loss of contralateral suppression (Froud et al., 2015), supporting the (Kim, 1986) theoretical postulate that the OHC – type II SGN afferent pathway contributes to the sensory drive of the MOC reflex. Conversely, in the same WT and *Prph*KO samples, there were ∼ 2.3 MOC efferent boutons per OHC for both genotypes (Froud et al., 2015), consistent with the retained olivocochlear efferent functionality shown here with electrical stimulation of the *Prph*KO COCB.

*In vivo* electrophysiological studies of type II SGN are limited, but suggest that these small, unmyelinated neurons have slow conduction velocities, high thresholds and relative insensitivity to sound stimulation. Robertson (1984) delivered horseradish peroxidase during intracellular high-impedance microelectrode recordings of SGN somata in the guinea pig Rosenthal’s canal to definitively label an outer spiral fibre projecting to basal OHCs. Unlike type I SGN, which project as radial fibres to IHCs and respond to sound stimulation, this single labelled type II SGN was ‘silent’ (Robertson, 1984). The more definitive data of Brown (1994) (Brown, 1994) is from 19 putative guinea pig type II units recorded with high impedance glass microelectrodes. These unlabelled, but presumed type II SGN, were characterised by longer antidromic response latencies than type I SGN units. Some responded to acoustic stimulation, with the most sensitive threshold at 80 dB SPL, matching the current contralateral suppression paradigm. Units with the longest antidromic latencies were generally unresponsive to sound. In a subsequent guinea pig study (Robertson et al., 1999), additional SGN recordings from “silent” units with long antidromic stimulus latencies were evaluated, but the identity and integrity of these cells was questioned.

Additional studies of type II SGN physiology have been performed *in vitro*. *In situ* whole-cell patch-clamp analysis in a rat neonatal cochlear slice preparation showed comparable recruitment of action potentials in type I and type II SGN, with an A-type inactivating K^+^ channel conductance prominent in the type II SGN, with reduced AMPA-type glutamate receptor conductance, but comparable ATP-activated inward currents (Jagger and Housley, 2003). Type II SGN had significantly depolarized resting potentials compared with type I SGN, which may explain the limited ability to record action potentials from these neurons *in vivo* with microelectrodes. Under current-clamp, isolated mouse type II SGN exhibited region-dependent variation in firing properties similar to those of type I SGN (Reid et al., 2004). Patch-clamp recordings of neonatal rat type II SGN terminal processes provided definitive evidence of OHC glutamatergic neurotransmission and purinergic neuromodulation, including spontaneous excitatory post-synaptic potentials/currents (EPSP/EPSC) (Weisz et al., 2009). Glutamatergic transmission appeared weaker than that of IHC – type I SGN synapses, based on the limited number of spontaneous EPSC events. These studies were extended and showed synaptic transmission via GluA2-containing AMPA receptors evoked by local application of K^+^. The data are consistent with a requirement for integrated summation of synaptic transmission from ∼ 6 OHCs to reach action potential threshold, sustained by an outer spiral fibre length constant beyond 1 mm, and supported by Na_v_1.6 voltage-gated Na^+^ channels (Weisz et al., 2012, Weisz et al., 2014, Martinez-Monedero et al., 2016).

There is overlapping distribution of type I and type II spiral ganglion afferent input to the cochlear nucleus. Both type I and type II SGN have branching termination in the dorsal and ventral cochlear nuclei, with type II SGN having considerable additional input to the cochlear nucleus granule cell region, a feature lacking in type I SGN (Brown et al., 1988a). In the small cell region in the posteroventral cochlear nucleus region (PVCN) immediately adjacent to the granule cell domain (Osen, 1969), electron microscopy localized type II synapses on stellate cells (Benson and Brown, 2004). This region also receives high-threshold type I auditory nerve fibres (Ryugo, 2008), has high-threshold auditory neurons (Ghoshal and Kim, 1997), and projects to the olivocochlear neurons (Ye et al., 2000). Darrow et al. (Darrow et al., 2012) used dye injections into the mouse cochlea for retrograde labelling of contralateral MOC efferent neurons in the ventral nucleus of the trapezoid body, alongside anterograde labelling from the PVCN that resolved planar stellate cells synapsing onto those MOC somata and dendrites; local PVCN ablation reduces MOC reflex suppression (Brown et al., 2003). Alongside this, studies across species have identified reciprocal connectivity between the MOC neurons and the cochlear nuclei, reflecting complexity of afferent and efferent circuitry regulating the MOC reflex (Robertson and Winter, 1988, Thompson and Thompson, 1991). A notable finding in the gerbil (Ryan et al., 1990) and mouse (Brown et al., 1988b) is that MOC efferent collaterals project to the cochlear nucleus granule cell region, a selective convergence with type II SGN afferent input. In support of the *in vivo* electrophysiological findings for type II SGN functionality, using noxious noise levels Flores *et al*. (Flores et al., 2015) reported increase in *cFos* signal in the granule cell region in *VGlut3* KO mice, which lack IHC – type I SGN transmission. This finding has been advanced by a more definitive study which selectively modulated OHC glutamatergic neurotransmission to demonstrate functional sensory drive by type II SGN afferents to the granule cell region of the cochlear nucleus spanning moderate (80 dB SPL) to damaging (115 dB SPL) sound levels (Weisz et al., 2021).

It is clear from the present study that the loss of acoustically-evoked contralateral suppression in the *Prph*KO mice occurs upstream of the superior olivary complex, in the afferent supply to the reflex, since the COCB remains functional. This is counter to the Maison et al. (Maison et al., 2016) *Prph*KO study, which did not observe the type II SGN morphological disruption, and where a negative result for electrically-evoked COCB suppression of the DPOAE in two *Prph*KO mice led to an inference that loss of contralateral suppression likely arose from failure of the efferent arm of the circuit. The present study verifies type II SGN synapse disruption and dendrite loss in the *Prph*KO mouse line. With the strong evidence (Weisz et al., 2021) that type II SGN are acoustically active at moderate and high sound levels, the loss of contralateral suppression in the *Prph*KO mice seems likely due to the type II SGN synapse disruption. A radical change in drive from the IHC – type I afferent sensory input in the *Prph*KO mouse appears less likely, since hearing sensitivity is maintained. However, other alternatives are possible. Peripherin is expressed centrally and brainstem disruptions could underlie the observed effects, although peripherin-immunopositive somata were not evident in the cochlear nucleus region (Barclay et al., 2007). Loss of peripherin-expressing interneuron connections from the cochlear nucleus to the MOC (if existent), or the small number of the MOC tunnel crossing fibres in mice that are peripherin immunopositive (Maison et al., 2016) are possibilities. In the cochlea, type II SGN fibres make terminal arborizations with Deiters’ and Hensen’s cells, particularly in the apical region, although it is uncertain whether these reflect synaptic couplings (Fechner et al., 2001). Reciprocal transmission between type II SGN afferents and the OHCs, evident from electron microscopy studies in cat and primates, could also contribute to local integration of OHC sensory transduction (Thiers et al., 2008). There are also synaptic varicosities between type II SGN afferents and the MOC efferent terminals (Ginzberg and Morest, 1984) which may support GABAergic transmission as part of a reciprocal microcircuit in the early post-natal period in rats (Weisz, 2020). Changes in these cochlear connections could potentially be a factor.

Any of these potential mechanisms demonstrate a critical role for peripherin in afferent feed to contralateral suppression. However, in the absence of evidence supporting alternatives, we feel that parsimony favors loss of OHC - type II SGN input as the root cause of loss of contralateral suppression. Parenthetically, a mouse study that selectively reduced IHC – type I afferent input by treating the cochlear round window membrane with ouabain, causing ∼ 90% reduction in the ABR wave I amplitude, showed no impact on contralateral suppression from 76 dB SPL broadband noise (Li et al., 2018). Given the long-standing evidence that type I SGN afferent input drives the MOC efferents, based on near-equivalency of thresholds and tuning characteristics (Robertson and Gummer, 1985, Liberman and Brown, 1986), the association here of disruption of type II SGN input and loss of contralateral suppression suggests that the broad reduction of the cochlear amplifier by the COCB at moderate to high sound levels captured by DPOAE may require the combined drive of both type I SGN and type II SGN input, with the unexpected finding that type I SGN drive alone may fall short. These findings provide a clear imperative for development of more sophisticated models for selective manipulation of these two cochlear afferent populations to probe the sensory drive of the MOC efferent control of the cochlear amplifier.

It is well established that MOC efferent feedback is critical to the long-term protection of the cochlear hair cells and their afferent innervation (reviewed by (Fuchs and Lauer, 2019)). MOC efferent – mediated protection from noise-induced hearing loss (NIHL) has been confirmed in mice using selective genetic manipulation of the OHC α9 / α10 nicotinic cholinergic receptor (nAChR). As noted above, a gain of function mutation of the α9 nAChR subunit confers heightened resistance to NIHL (Taranda et al., 2009), while an α9 knockout exhibited enhanced NIHL (Maison et al., 2002). Moreover, chronic moderate (84 dB SPL) noise exposure caused increased type I SGN synaptopathy in mice when the COCB was sectioned at the floor of the fourth ventricle (Maison et al., 2013). Here, in the *Prph*KO mouse model, we demonstrate clear association between the disruption of OHC – type II SGN sensory input, near-elimination of MOC efferent – mediated contralateral suppression at moderate to high sound levels, and reduction in otoprotection against NIHL.

In summary, these findings suggest that from moderately loud sound levels, the cochlear OHC-type II SGN afferent pathway contributes to MOC efferent regulation of the cochlear amplifier that confers protection from noise-induced hearing loss.

## Acknowledgements

This work was supported by Australia grants from the National Health & Medical Research Council (NHMRC) APP1052463, APP1189113, APP1188643 & U.S. Department of Veterans Affairs grants BX001205 and RX002704. Funding was also provided by a donation from Alan and Lynne Rydge. Edward Crawford is thanked for his contributions to development of the instrumentation and software for the study.

---

**Figure 2-figure supplement 1:**
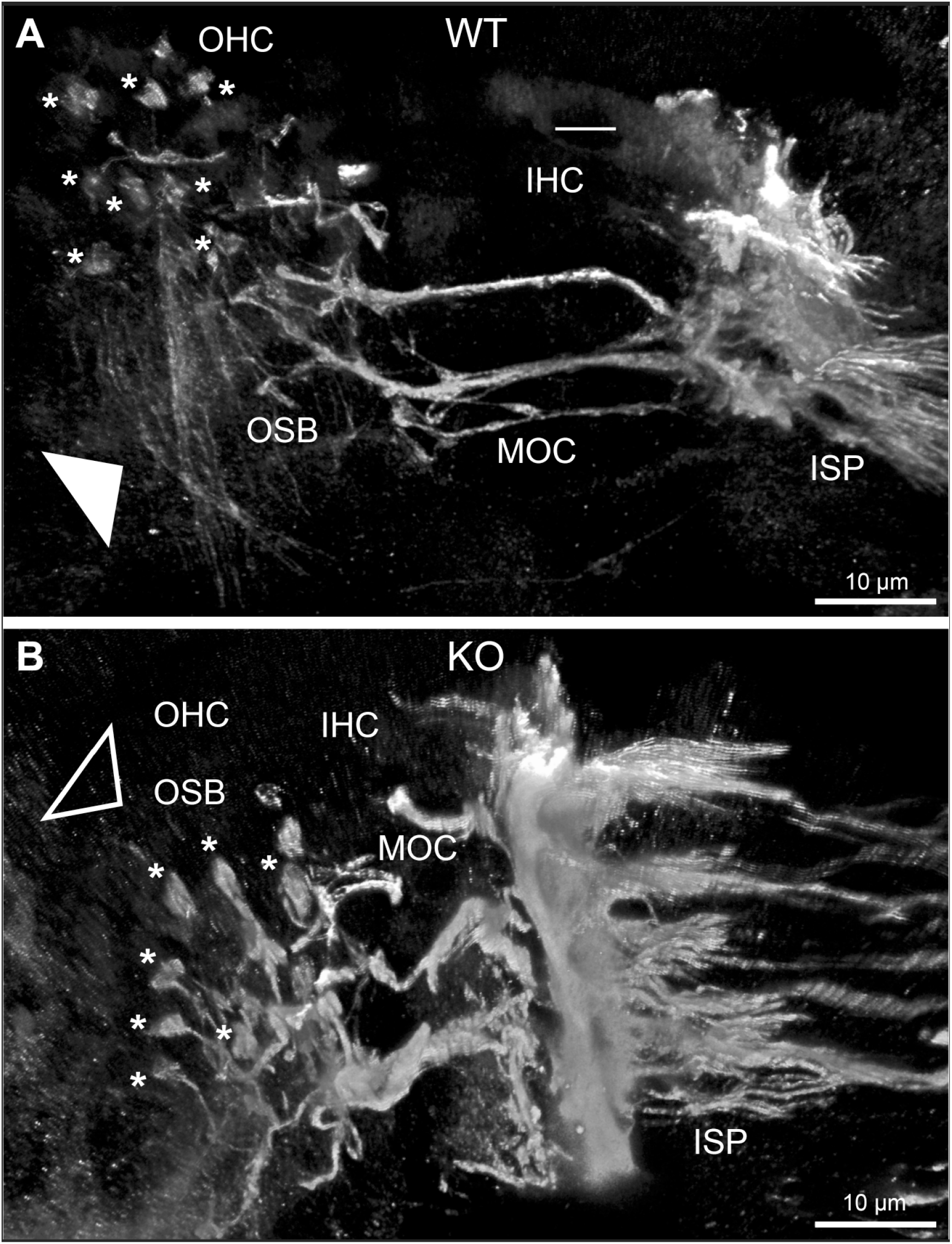
Disruption of outer spiral bundle structure (OSB; type II spiral ganglion neuron dendrites (type II SGN)) in mid -cochlear region of the Prph knockout (KO) resolved using β-III tubulin immunofluorescence. Confocal transparent-mode 3D reconstructed images of batch-processed WT and PrphKO cryosections with the basilar membrane surface oriented uppermost to optimally delineate the OSB. **A**, In the WT cochlea, the outer spiral fibres (type II SGN) within the OSB (arrowhead) run vertically below the three rows of outer hair cells (OHCs). The medial olivocochlear (MOC) efferent fibers project horizontally from the inner spiral plexus (ISP) region, to the OHC region, where they branch to form large synaptic boutons (*) at the base of multiple OHCs. **B**, In the PrphKO image, the OSB are not evident (open arrowhead), while equivalent MOC efferent fiber projections extend to the OHCs and form synaptic boutons (*). IHC, inner hair cells. See also Figure 2.

**Figure 4-figure supplement 1:**
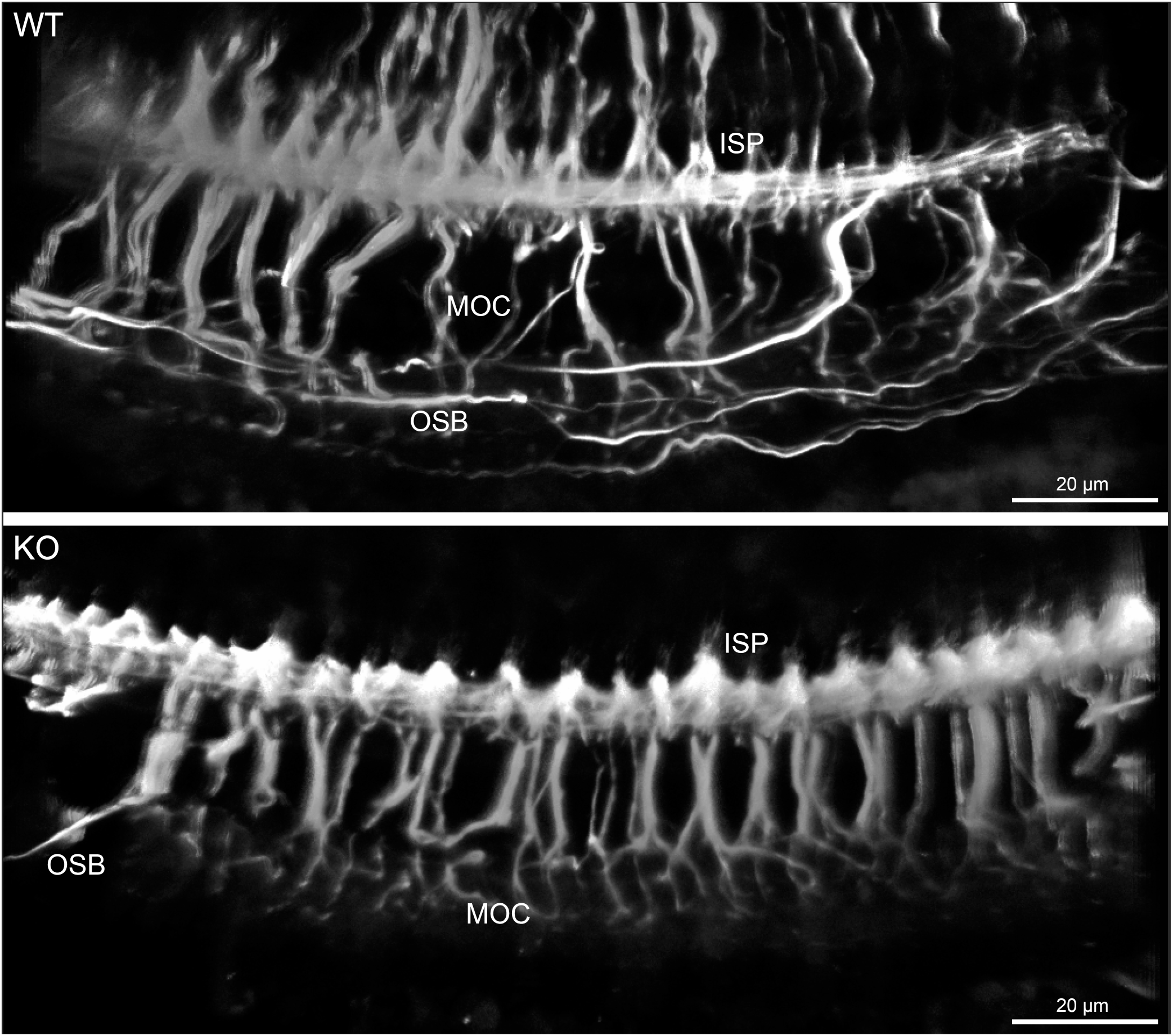
NF200 immunofluorescence labelling (maximum intensity projections) of batch-processed wholemounts from the apical region of wildtype (WT) and PrphKO organ of Corti. Note the lack of outer spiral fibres within the outer spiral bundle (OSB; type II SGN afferent fibres) in the KO tissue, while medial olivocochlear (MOC) efferent projections to the outer hair cell region from the inner spiral plexus region (ISP) are equivalent. To optimally resolve the OSB fibre tracts, the tissue was imaged with the basilar membrane surface closest to the microscope objective, as the OSB lies beneath the MOC fibres. See also Figure 4.

**Figure 6-figure supplement 1:**
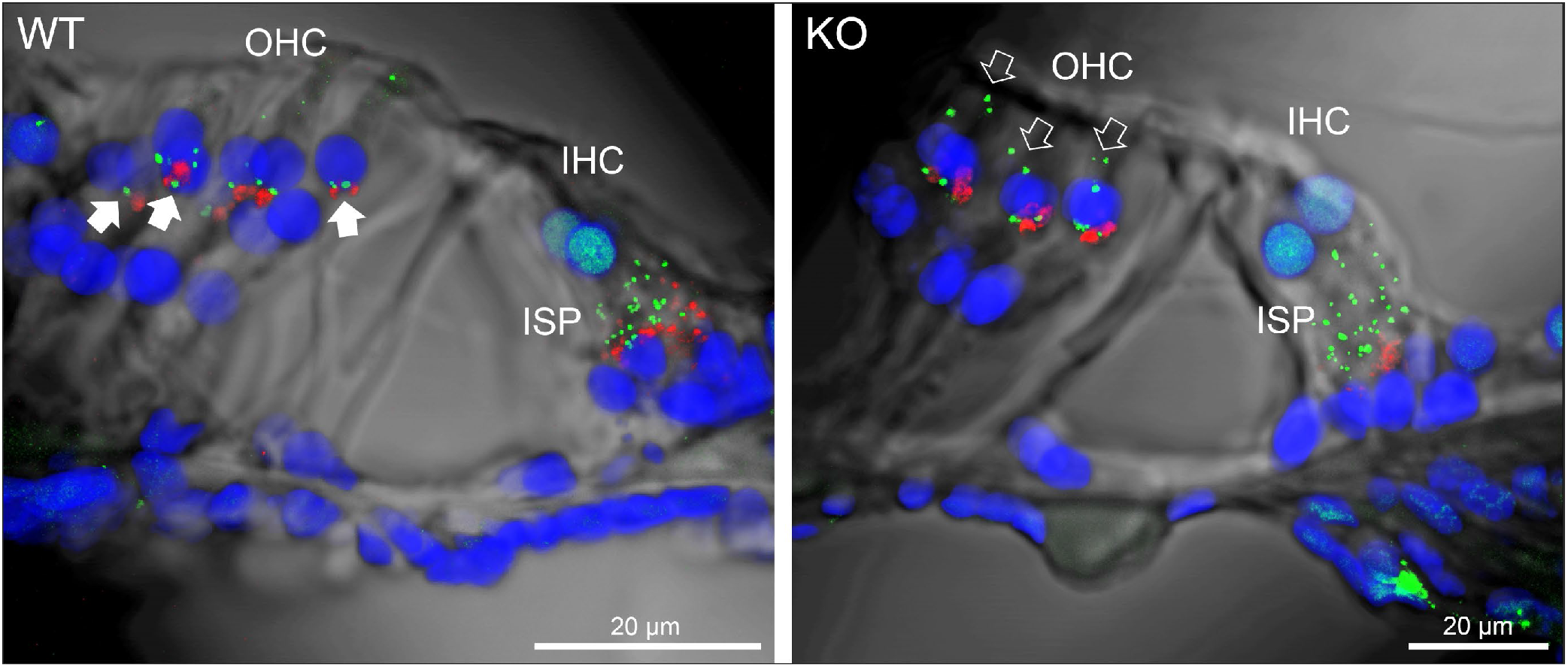
Evidence of disruption of the type II SGN afferent synapses with the outer hair cells (OHC) in the PrphKO cochlea. CtBP2/RIBEYE immunofluorescence (green puncta) localizes pre-synaptic ribbon complexes at the base of OHCs and inner hair cells (IHC). While this labelling is consistent between WT and KO at the IHC synapses (inner spiral plexus region, ISP), there is pronounced apical migration of many of the synaptic ribbons away from the synaptic pole of the KO OHCs (compare filled arrows for WT and open arrows for KO). In contrast, the medial olivocochlear efferent synaptic boutons (VAChT immunofluorescence, red) are retained in the mid and medial aspects of the synaptic pole of the OHCs in the WT and KO cochlea. Nuclei labelled with DAPI. 50 µm cryosections. See also Figure 6.

**Figure 9-figure supplement 1.**
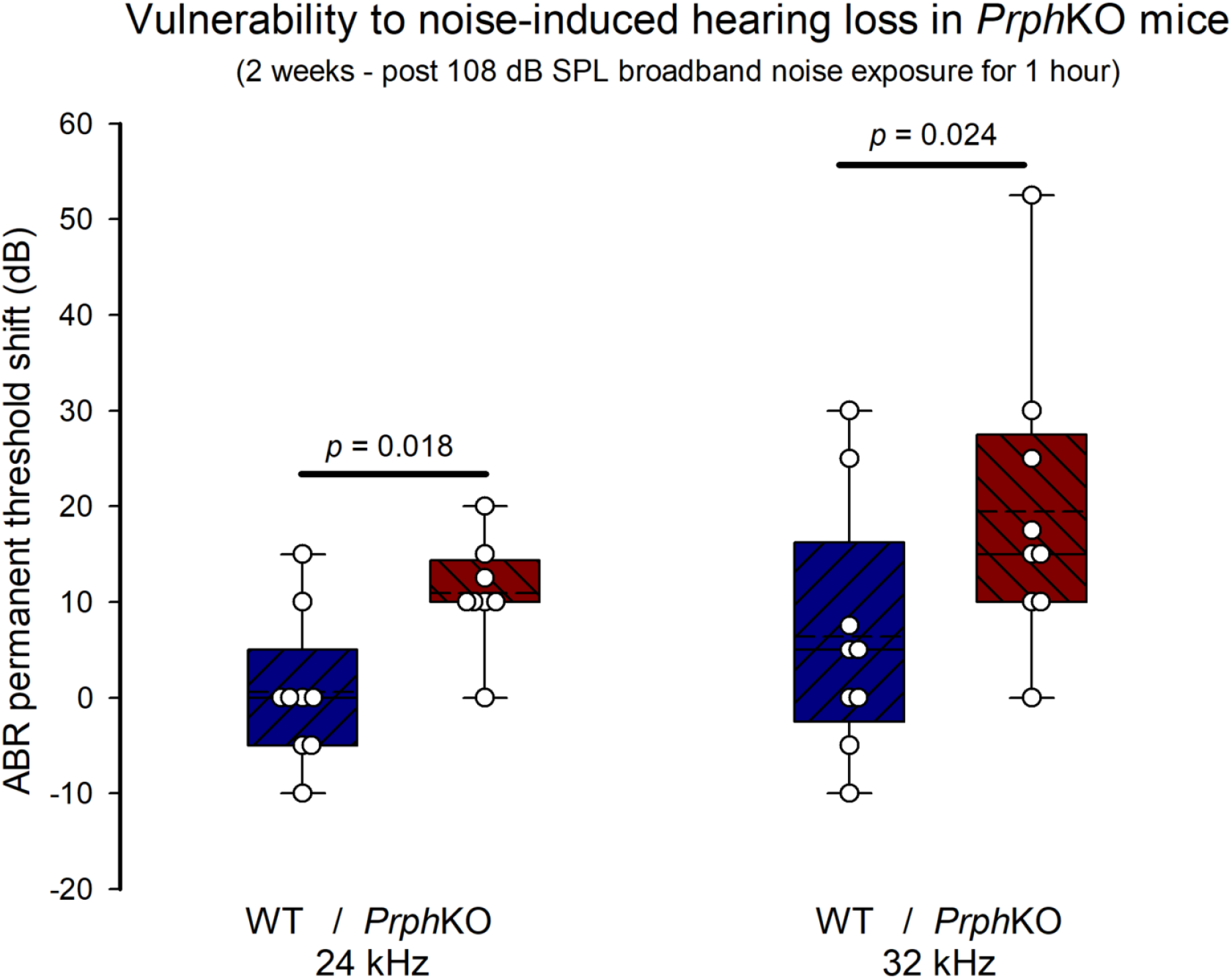
ABR threshold shifts indicative of increased vulnerability to noise-induced hearing loss in PrphKO. Boxplots show the 25% and 75 % boundaries and 95% limits along with overlaid data from 9 WT and 9 PrphKO mice; dashed line indicates mean, solid line indicates median. (Holm-Sidak post-hoc comparisons, repeated measures ANOVA; Grubbs’ test (GraphPad) excluded one data point as an outlier for PrphKO 24 kHz)). See also Figure 9; Figure 9-Source Data 1.

**Figure 9-figure supplement 2.**
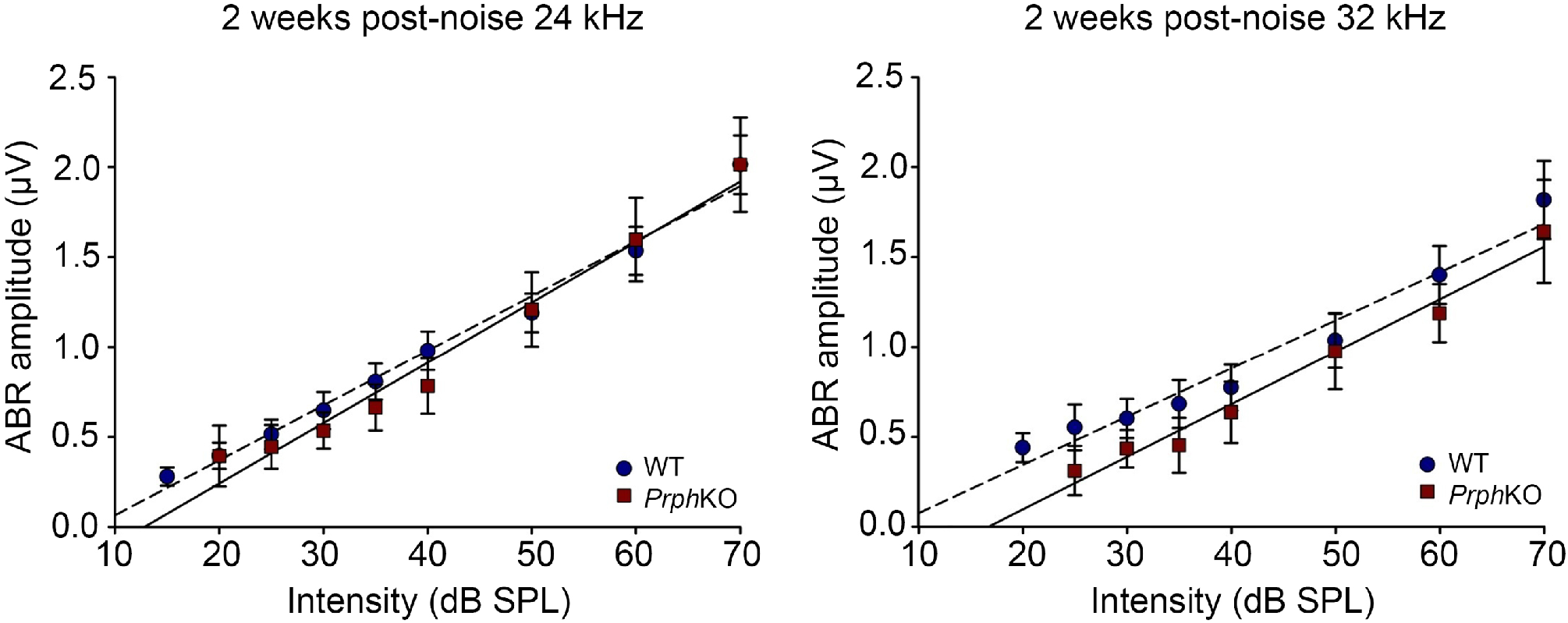
Suprathreshold ABR wave N1 – P2 amplitudes reflect neural recruitment with increasing sound intensity two weeks after 108 dB SPL white noise exposure in WT mice, and the PrphKO mice with significant hearing loss. The slopes of the linear regression best fits to these input / output functions at 24 kHz and 32 kHz were not significantly different across genotype (n = 8 WT; n = 7 PrphKO – solid lines). Average slopes & R^2^ values: 24 kHz: WT 0.030 µV / dB SPL & 0.988; KO 0.033 µV / dB SPL & 0.975; 32 kHz: WT 0.026 µV / dB SPL & 0.962; KO 0.029 µV / dB SPL & 0.98). See also Figure 9.

**Figure 6-Source Data 1.**
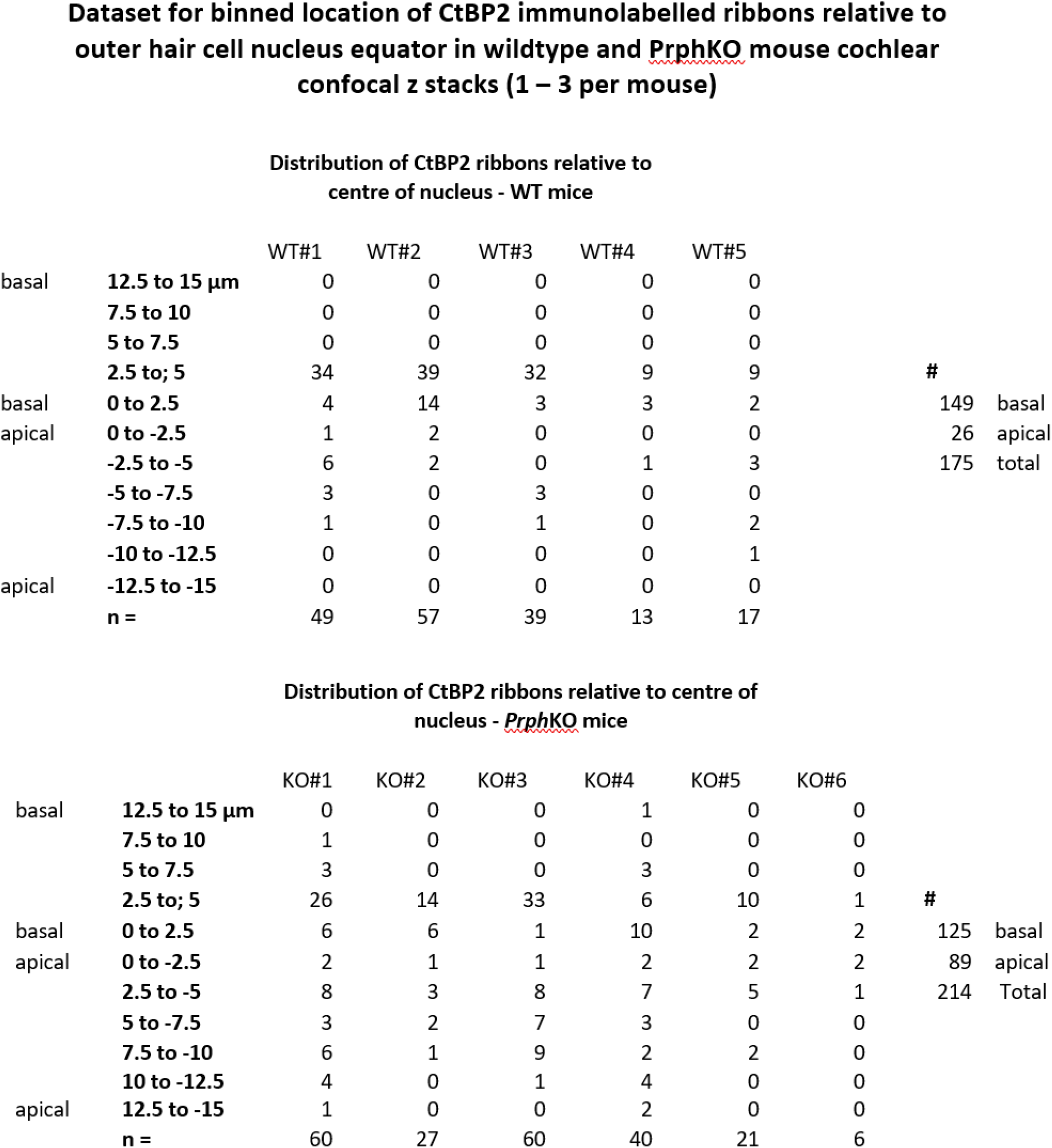

**Figure 9-Source Data 1.**
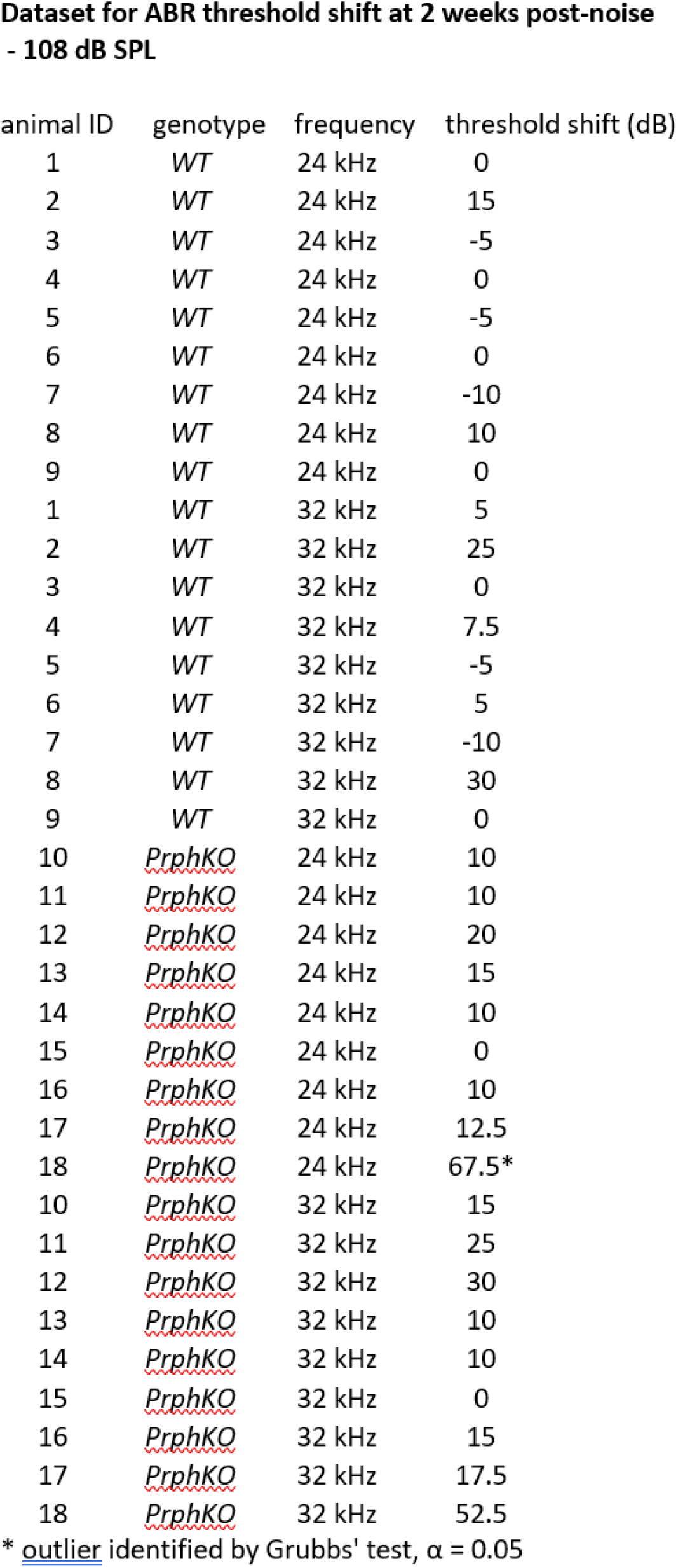

